# Na^+^/K^+^ ATPase-Ca_v_1.2 nanodomain differentially regulates intracellular [Na^+^], [Ca^2+^] and local adrenergic signaling in cardiac myocytes

**DOI:** 10.1101/2023.08.31.553598

**Authors:** Mariusz Karbowski, Liron Boyman, Libet Garber, Humberto C. Joca, Nicolas Verhoeven, Andrew K. Coleman, Christopher W. Ward, W. Jonathan Lederer, Maura Greiser

## Abstract

**Background:** The intracellular Na^+^ concentration ([Na^+^]_i_) is a crucial but understudied regulator of cardiac myocyte function. The Na^+^/K^+^ ATPase (NKA) controls the steady-state [Na^+^]_i_ and thereby determines the set-point for intracellular Ca^2+^. Here, we investigate the nanoscopic organization and local adrenergic regulation of the NKA macromolecular complex and how it differentially regulates the intracellular Na^+^ and Ca^2+^ homeostases in atrial and ventricular myocytes.

**Methods:** Multicolor STORM super-resolution microscopy, Western Blot analyses, and in vivo examination of adrenergic regulation are employed to examine the organization and function of Na^+^ nanodomains in cardiac myocytes. Quantitative fluorescence microscopy at high spatiotemporal resolution is used in conjunction with cellular electrophysiology to investigate intracellular Na^+^ homeostasis in atrial and ventricular myocytes.

**Results:** The NKAα1 (NKAα1) and the L-type Ca^2+^-channel (Ca_v_1.2) form a nanodomain with a center-to center distance of ∼65 nm in both ventricular and atrial myocytes. NKAα1 protein expression levels are ∼3 fold higher in atria compared to ventricle. 100% higher atrial I_NKA_, produced by large NKA “superclusters”, underlies the substantially lower Na^+^concentration in atrial myocytes compared to the benchmark values set in ventricular myocytes. The NKA’s regulatory protein phospholemman (PLM) has similar expression levels across atria and ventricle resulting in a much lower PLM/NKAα1 ratio for atrial compared to ventricular tissue. In addition, a huge PLM phosphorylation reserve in atrial tissue produces a high ß-adrenergic sensitivity of I_NKA_ in atrial myocytes. ß-adrenergic regulation of I_NKA_ is locally mediated in the NKAα1-Ca_v_1.2 nanodomain via A-kinase anchoring proteins.

**Conclusions:** NKAα1, Ca_v_1.2 and their accessory proteins form a structural and regulatory nanodomain at the cardiac dyad. The tissue-specific composition and local adrenergic regulation of this “signaling cloud” is a main regulator of the distinct global intracellular Na^+^ and Ca^2+^ concentrations in atrial and ventricular myocytes.

## Introduction

An important but often overlooked ’silent’ partner of cardiac myocyte function is the intracellular Na^+^ concentration ([Na^+^]_i_). [Na^+^]_i_ is widely acknowledged as a crucial regulator of the intracellular Ca^2+^ concentration ([Ca^2+^]_i_) in the cytosol, the sarcoplasmic reticulum (SR), and the mitochondria through the richly abundant Na^+^/Ca^2+^ exchanger (NCX) in the plasma membrane and the mitochondrial Na^+^/Ca^2+^ exchanger (NCLX)^1–4^, which exchange 1 Ca^2+^ ion for 3 Na^+^ ions ^4–7^. The pivotal role played by [Na^+^]_i_ becomes evident when its regulation is disrupted, for example in heart failure, where a significant increase in the [Na^+^]_i_ in ventricular myocytes is a major contributing factor in the pathological elevation of [Ca^2+^]_i_, which is a hallmark of heart failure^8, 9^. High [Na^+^]_i_ and [Ca^2+^]_i_ are key contributors to heart failure progression and underlie the development of malignant cardiac arrhythmias^10, 11^. Similarly, changes in [Na^+^]_i_ in atrial myocytes, possibly due to an increased influx of Na^+^ into the cell, are thought to contribute to the development of atrial fibrillation^12, 13^.

Previous work by Mohler and others showed that Na^+^ transport proteins co-cluster on the sarcolemmal membrane and the transverse tubular system in cardiac myocytes. Specifically, the Na^+^/K^+^ ATPase (NKA), the only major Na^+^ extrusion transporter in all cells, and the NCX co-localize in an ion channel complex tethered by the structural protein ankyrin-B (ankB)^14^. Genetic deficiency of ankB, which disrupts the nanostructure of the Na^+^ transporter cluster, leads to atrial and ventricular arrhythmias, likely by disrupting the intracellular Ca^2+^ and Na^+^ homeostasis^15, 16^. The results of these studies indicated that the nanoscopic spatial organization of Na^+^ transporters appeared to be crucial for the regulation of intracellular Na^+^ and Ca^2+^ signaling. Historically, inhibition of the NKA by cardiotonic steroids was used to increase cardiac inotropy by exploiting the tight regulatory connection between [Na^+^]_i_ and [Ca^2+^]_i_. Inhibition of the NKA, which extrudes 3 Na^+^ ions and imports 2 K^+^ ions, increases [Na^+^]_i_ and subsequently [Ca^2+^]_i_ by decreasing the NCX’s driving force for Ca^2+^ efflux. Nanoscopic composition, size and regulation of Na^+^ transport hubs and how they might be linked, structurally or functionally, to other ion transport domains remains poorly understood.

A crucially important intracellular Ca^2+^ signaling hub, the cardiac dyad, is much better characterized. It consists of a small number of L-type Ca^2+^ channels (Ca_v_1.2) that serve as sarcolemmal voltage-sensors in close apposition to an intracellular calcium-release mechanism consisting of a neighboring cluster of ryanodine receptors (RyR2) 20 nm away^17^. This close proximity enables the opening of Ca_v_1.2 to trigger Ca^2+^ sparks from the apposing RyR2 clusters by Ca^2+^-induced Ca^2+^-release (CICR) ^18^. An early concept of functional interaction between Na^+^ transport proteins and the cardiac dyad suggested that a functional coupling of the NCX to the L-type Ca^2+^ channel in a subsarcolemmal “fuzzy space” might regulate [Ca^2+^]_i_ and [Na^+^]_i_ in cardiac myocytes^3^. Recent technical advances in multi-color super-resolution microscopy have opened a new track for high resolution quantitative investigations into the nanoscopic spatial organization of Na^+^ and Ca^2+^ transport hubs and their potential structural and functional linkage, which may prove critical for our understanding of [Na^+^]_i_ and [Ca^2+^]_i_ regulation in cardiac myocytes. Specifically, an xy resolution ∼30 nm, which can only be achieved by true super-resolution microscopy techniques like STORM and STED microscopy, is required to investigate nanodomain formation of cardiac ion channels.

In the present study, using two-color STORM super-resolution microscopy, we made the surprising discovery that NKA (NKAα1) tightly co-clusters with the L-type Ca^2+^ channel (Ca_V_1.2) at the cardiac dyad. We performed a quantitative and qualitative investigation of this newly discovered NKAα1-Ca_V_1.2 nanodomain and how its tissue-specific composition and function underlies differences in global [Na^+^]_i_ and its regulation in atria and ventricles. Close nanoscopic alignment of NKAα1 and Ca_V_1.2 clusters (center-to-center distance ∼65 nm) creates a shared signaling domain between the two ion channels and their associated regulatory proteins. Specifically, functional studies showed that this shared “signaling cloud” provides local, PKA-mediated regulation of the NKA current (I_NKA_) through A-kinase anchoring proteins (AKAPs). Chamber-specific analysis showed that NKAα1 has ∼3fold higher expression levels in atrial versus ventricular tissue. NKAα1 aggregates in large clusters on the cell membrane and the membranes of the transverse or axial tubule system (TATS) in atrial myocytes. These NKA “superclusters” generate a much larger I_NKA_ in atrial myocytes than in ventricular myocytes, where NKA density is much lower. In addition, the NKA’s Na^+^ affinity is much higher in atrial tissue than in ventricle. Phospholemman (PLM), the NKA’s regulatory protein, has similar expression levels across atrial and ventricular tissue. Given the high abundance of NKAα1 in atria, the PLM/NKAα1 ratio is much lower in atrial compared to ventricular tissue. PLM, when associated with NKAα1, decreases the NKA’s Na^+^ affinity. In addition, functional and in vivo biochemical data show that atrial NKAα1 superclusters are highly sensitive to ß-adrenergic activation due to a very high phosphorylation capacity of PLM in atria.

Taken together, the NKAα1-Ca_V_1.2 complex forms a newly discovered Na^+^-Ca^2+^ nanodomain with a shared “signaling cloud” that is part of the cardiac dyad. The atrial specific composition of the NKAα1-Ca_V_1.2 signaling cloud provides robust regulation of atrial Na^+^ homeostasis with an increased margin of safety for a system with known arrhythmogenic vulnerability.

## Methods

A detailed description of all experimental procedures and statistical tests can be found in the Supplemental Methods.

## Results

The advancement of multi-color super-resolution microscopy techniques has enabled novel investigations into structural and functional nanodomains in cardiac myocytes. Here, we used two-color direct STORM super-resolution imaging to investigate the structural composition of Na^+^ signaling nanodomains and how they may be linked to the cardiac dyad as a starting point into an investigation into local regulation of the intracellular Na^+^ homeostasis. We discovered that Na^+^/K^+^-ATPase-α1 (NKAα1) and the pore-forming subunit of the L-type Ca^2+^ channel (Ca_v_1.2) form a tightly co-clustered nanodomain at the cardiac dyad. The association of the only major Na^+^ extrusion transporter with the major (voltage-gated) Ca^2+^ influx channel was surprising and had not been conceptualized previously. We then investigated whether quantitative and qualitative differences in the composition of this newly discovered nanodomain give rise to differences in [Na^+^]_i_ and its regulation in atrial and ventricular myocytes.

*The cardiac NKAα1-Ca_v_1.2 nanodomain.* Using two-color STORM super-resolution microscopy, we found that NKAα1, the dominant cardiac isoform of the NKAα subunit, co-clusters closely with the pore forming α1_C_ subunit of the L-type Ca^2+^ channel (Ca_v_1.2) in ventricular cardiac myocytes. Fig. 1A shows an exemplar of a two-color STORM image showing NKAα1 clusters (green) and Ca_v_1.2 clusters (red). Fig. 1B shows the image depicted in Fig.1A with masks generated for quantitative analysis of the ion cluster areas (NKAα1 = magenta, Ca_v_1.2 = turquoise). Fig. 1C shows, in blue, the shared area between NKAα1 and Ca_v_1.2 clusters. Quantitative examination revealed that the centers of NKAα1 and Ca_v_1.2 clusters are very close together (center-to-center distance 63 ± 3 nm, Fig. 1D). These data provide the first direct evidence that NKAα1 and Ca_v_1.2 form a novel structural nanodomain in cardiac myocytes. We next evaluated (1) whether the NKAα1 and Ca_v_1.2 nanodomain also exists in atrial myocardium and (2) whether its composition is qualitatively or quantitatively different in atrial compared to ventricular tissue. Fig. 2A-C shows an exemplar of a two-color STORM image in an isolated atrial myocyte depicting the co-clustering of NKAα1 (green) and Ca_v_1.2 (red) with masks generated for quantitative analysis (NKAα1 = magenta, Ca_v_1.2 = turquoise, Figs 2B and 2C). NKAα1-Ca_v_1.2 cluster center-to-center distance was similar in atrial myocytes (69 ± 2 nm, Fig. 2D) compared to the previously measured distance in ventricular myocytes. In atrial myocytes, 79% of all Ca_v_1.2 cluster area overlapped with NKAα1 cluster area (Fig. 2E). Surprisingly, atrial Ca_v_1.2 clusters were significantly larger than in ventricular myocytes (Fig. 2F).

**Figure 1.**
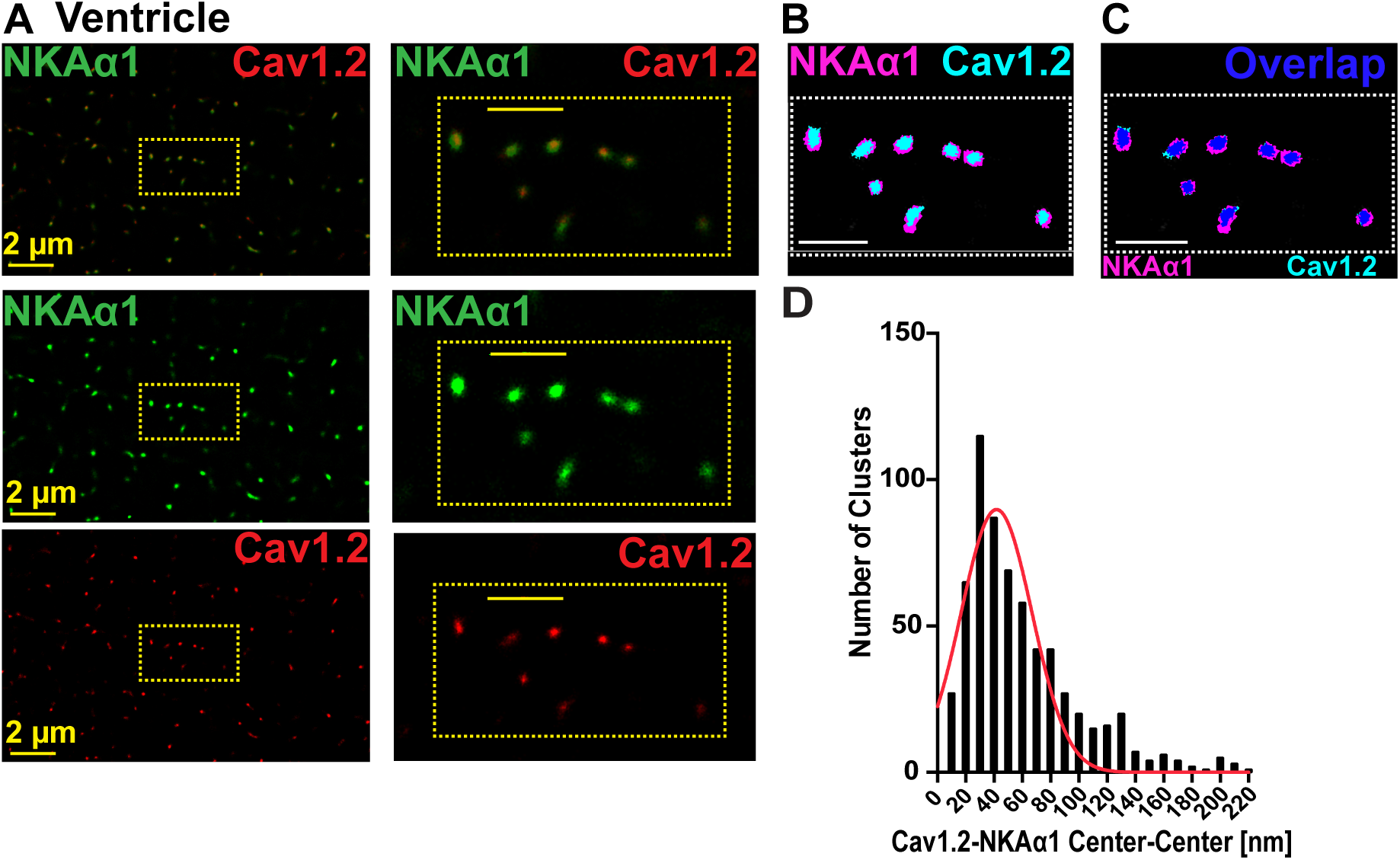
NKAα1 and Ca_v_1.2 nanodomain in ventricular myocytes. **A.** Two-color STORM image of NKAα1 and Ca_v_1.2 in a ventricular cardiac myocyte (Scale bar: 1 μm). **B.** Image as in A. Colored areas show masks generated for analysis. **C.** Image as in **B.** with the addition of the overlap area of NKAα1 and Ca_v_1.2 clusters (shown in blue). **D.** Center-to-center distance of NKAα1 and Ca_v_1.2 clusters (N = 3 animals, n = 5 cells).

**Figure 2.**
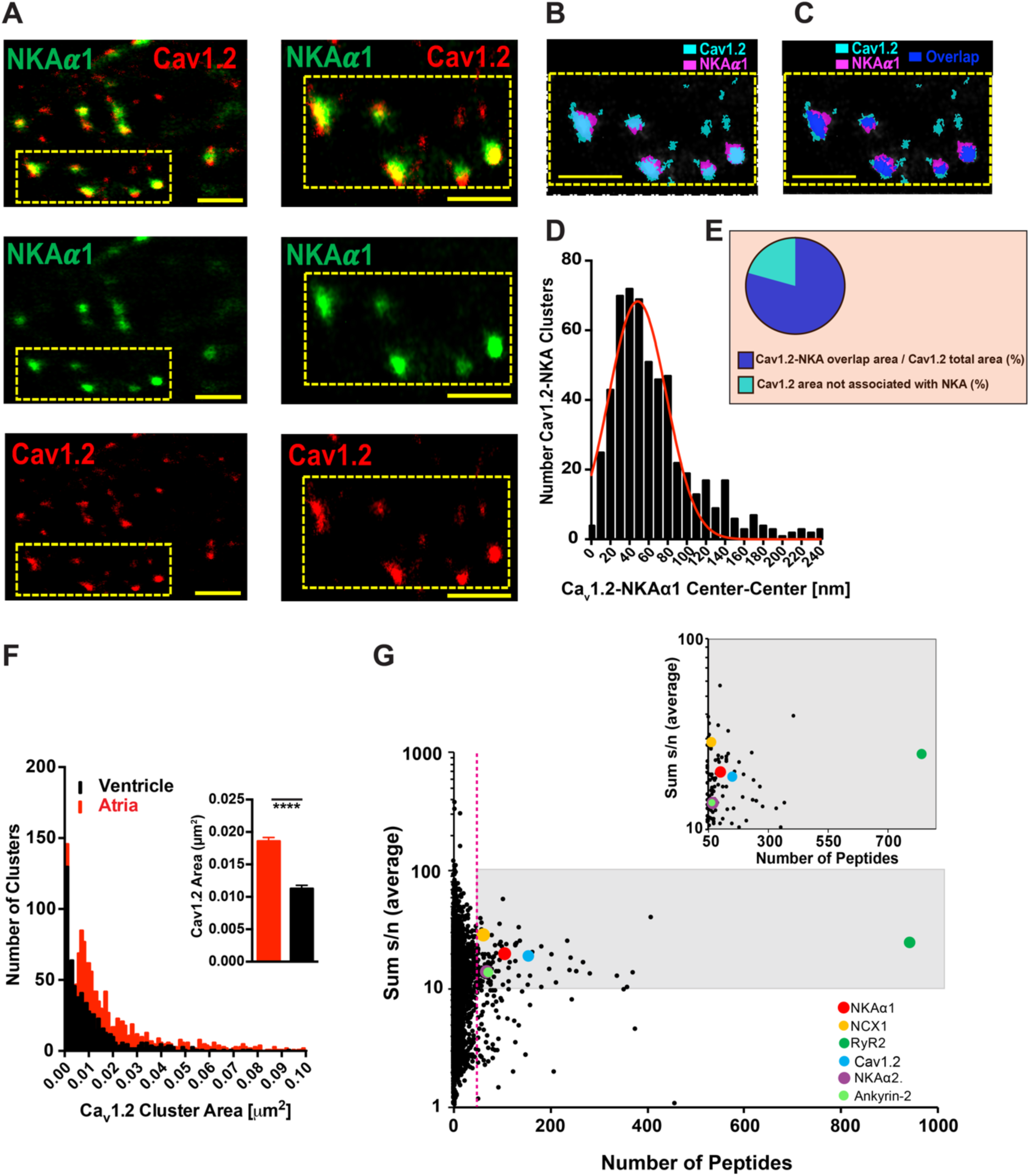
NKAα1 and Ca_v_1.2 signaling cloud in atrial myocytes. Two-color STORM image of NKAα1 and Ca_v_1.2 in an atrial cardiac myocyte (Scale bar: 1 μm). **B.** Image as in A. Colored areas show masks generated for analysis. **C.** Image as in **B.** with the addition of the overlap area of NKAα1 and Ca_v_1.2 clusters (shown in blue). **D.** Center-to-center distance of NKAα1 and Ca_v_1.2 clusters (N = 3 animals, n = 12 cells). **E.** Ca_v_1.2/NKAα1 overlap area (%). **F.** Ca_v_1.2 cluster size (N = 3 animals, n = 12 cells). **F.** Atrial Ca_v_1.2 cluster sizes. **G.** Proximity proteomics previously performed in mouse α1_C_-APEX (Ca_v_1.2) myocytes^19^. The number of peptides of each protein is plotted against the sum of its increase in α1_C_-APEX-labeled versus unlabeled myocytes. Dotted line depicts 50 peptides, which represent the top 2% out of all identified peptides. Inset shows identified proteins in the top 2% group (50 peptides) with more than 10fold increase in the α1_C_-APEX-labeled versus unlabeled myocytes (NCX1: Na^+^/Ca^2+^ exchanger, RyR2: ryanodine receptor type 2).

NKAα1-Ca_v_1.2 co-clustering was further corroborated by proximity proteomics previously performed by Marx and colleagues in mouse α1_C_-APEX (Ca_v_1.2) cardiac myocytes^19^. Replotting of these data shows that NKAα1 was in the top 2 % (1.36%) of peptide abundance within all α1_C_-APEX targets. This represented a ∼ 20fold increase in peptide abundance over non-APEX-labeled myocytes (Fig 2G). In addition, as shown in Fig 2G, NKAα2, NCX and Ankyrin-2 are also within the top 2% of peptide abundance in α1_C_-APEX myocytes.

We evaluated NKAα1 distribution in atrial and ventricular cardiac myocytes and performed quantitative analysis of NKAα1 cluster size using STORM super-resolution imaging. Figs. 3A and 3B show the distribution of NKAα1 (red) compared to a membrane marker, caveolin-3 (Cav3, green) in an atrial and a ventricular myocyte using conventional confocal imaging. Fig. 3C shows an exemplar of NKAα1 dSTORM imaging in an atrial myocyte. The upper panel shows the overview of the mouse atrial myocyte and the lower panels reveal the organization and large size of individual NKAα1 clusters at high resolution. Similarly, Fig. 3D depicts NKAα1 STORM imaging in a single ventricular myocyte. Analysis of NKAα1 cluster area reveals that NKAα1 clusters are substantially larger in atrial than in ventricular myocytes (Fig 3E). Figs 3F and 3G show the NKAα1 cluster size distribution for the cells shown in Figs 3C and 3D, respectively.

**Figure 3.**
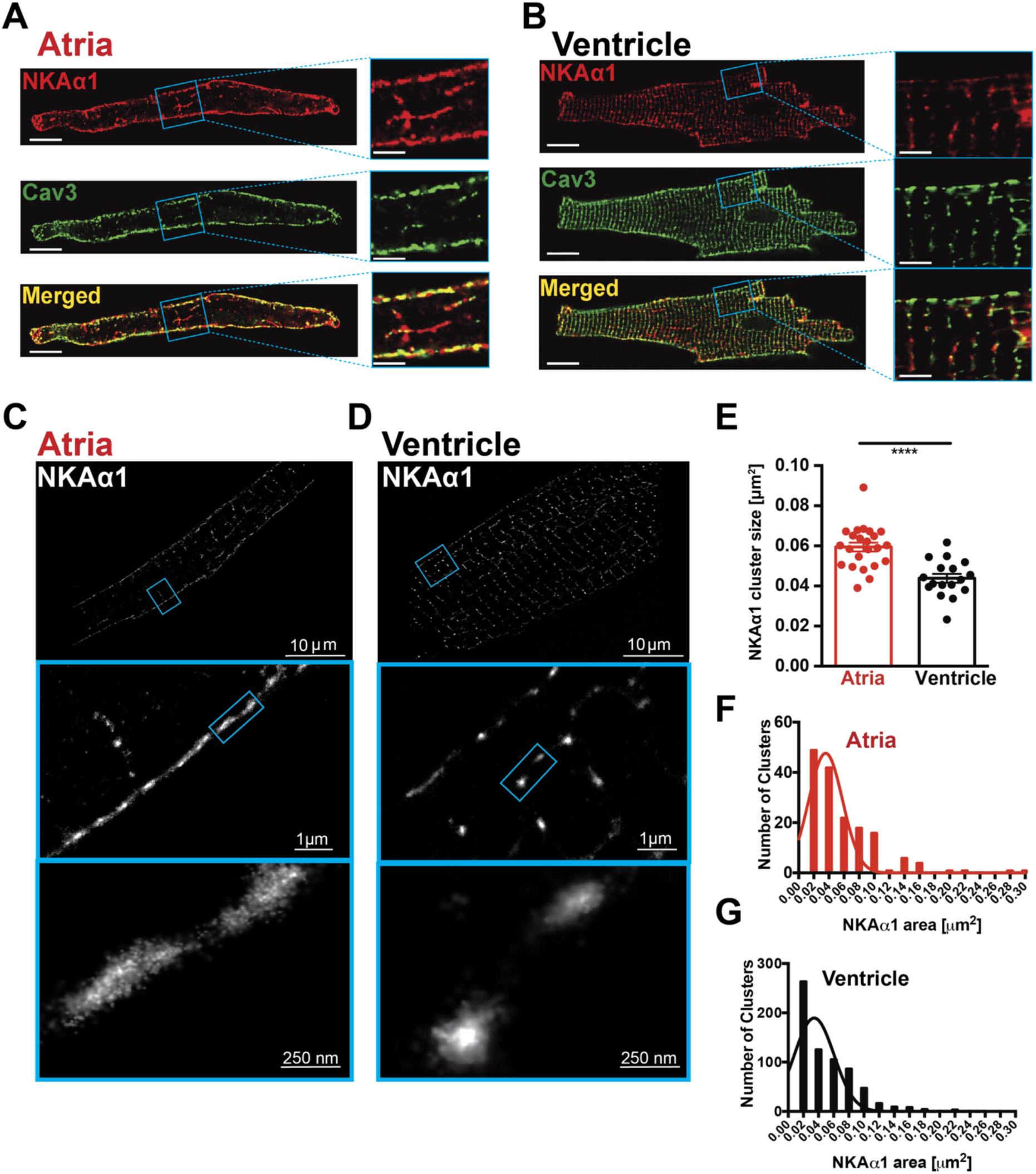
Super-resolution imaging of NKAα1 with dSTORM. **F.** Confocal image of an atrial myocyte stained with NKAα1 and Caveolin 3 (Cav 3). **G.** Confocal image of a ventricular myocyte stained with NKAα1 and Cav 3. **C.** Representative dSTORM image of NKAα1 clusters in an atrial myocyte (left panels) and in a ventricular myocyte (right panels). **D.** Mean NKAα1 cluster sizes in atrial and ventricular myocytes (4 animals; n = 23 atrial myocytes, n = 17 ventricular myocytes). **D.** Histograms showing NKAα1 cluster size distribution derived from **C**.

Taken together, super-resolution microscopy reveals that the NKAα1-Ca_v_1.2 structural nanodomain is present in both atrial and ventricular myocytes, suggesting that it is a preserved structural element across cardiac cells. Moreover, high resolution quantification of ion cluster areas provides direct evidence of tissue-specific composition of the NKAα1-Ca_v_1.2 nanodomain where atrial NKAα1 and Ca_v_1.2 form much larger “superclusters” than their ventricular counterparts.

### Tissue specific differences in the Na^+^/K^+^ ATPase (NKA) in heart

We next investigated how the tissue-specific composition of the NKAα1-Ca_v_1.2 nanodomain affects tissue-specific Na^+^ homeostasis and Na^+^ transport. NKA current (I_NKA_) was measured using the whole cell patch clamp technique. For the experiments shown in Fig. 4A, the cell interior is filled with 12 mM [Na^+^] from the patch-clamp pipette. At the beginning of the experiment, the NKA is completely blocked by the absence of extracellular K^+^. When the extracellular solution [K^+^] is increased from 0 mM to 5.4 mM, as is shown in the protocol line (green), the Na^+^ pump is activated and produces an outward I_NKA_. Fig. 4A shows the increase in outward I_NKA_ current normalized to the surface area (pA/pF) in an atrial (red) and a ventricular (black) myocyte. Fig. 4B depicts the distribution of I_NKA_ for multiple atrial and ventricular myocytes. The mean current density of I_NKA_ in atrial cells (red) is roughly twice as great as in ventricular myocytes (black). Furthermore, western blot analyses shown in Fig. 4C reveal that the NKAα1 isoform was roughly three times more abundant in atrial versus ventricular tissue while NKAα2 protein expression levels were similar in both tissues, as were protein expression levels of NCX. Given the already roughly 4-fold greater abundance of α1 over α2 isoforms in ventricles^20, 21^, the results indicate that the α1 isoform of NKA is even more dominant in atria than in ventricles. We further evaluated the functional contribution of NKAα1 and -α2 to total atrial I_NKA_ by measuring the concentration-dependence of ouabain-mediated inhibition of atrial I_NKA_. NKAα isoforms contain a highly conserved binding site for ouabain, a cardiotonic steroid, which is an NKA inhibitor^22, 23^. In rodents, cardiac NKAα1 and -α2 isoforms can be functionally distinguished by their different ouabain affinity. NKAα1 has low ouabain affinity (EC_50_ >> 1 μM) whereas NKAα2 has a high ouabain affinity in the low nmol/L range^21, 24^. Mean dose-response curves of atrial I_NKA_ inhibition are shown in Fig. 4D, measured with 12 mM [Na^+^] in the patch-clamp pipette. Data were best fit with a one-binding site equation (Fig. 4E), which indicates the presence of just one ouabain binding site. This low-affinity binding site (K_D_ = 83.40 ± 8.55 μM) can be attributed to NKAα1^21^. The lack of a second, high-affinity ouabain binding site, which was previously shown in ventricular myocytes^21^, suggests that NKAα1 is so dominantly expressed in atrial myocytes, as shown in the Western Blot data, that the contribution of NKAα2, the putative high-affinity ouabain binding site, to atrial I_NKA_ is negligible.

**Figure 4:**
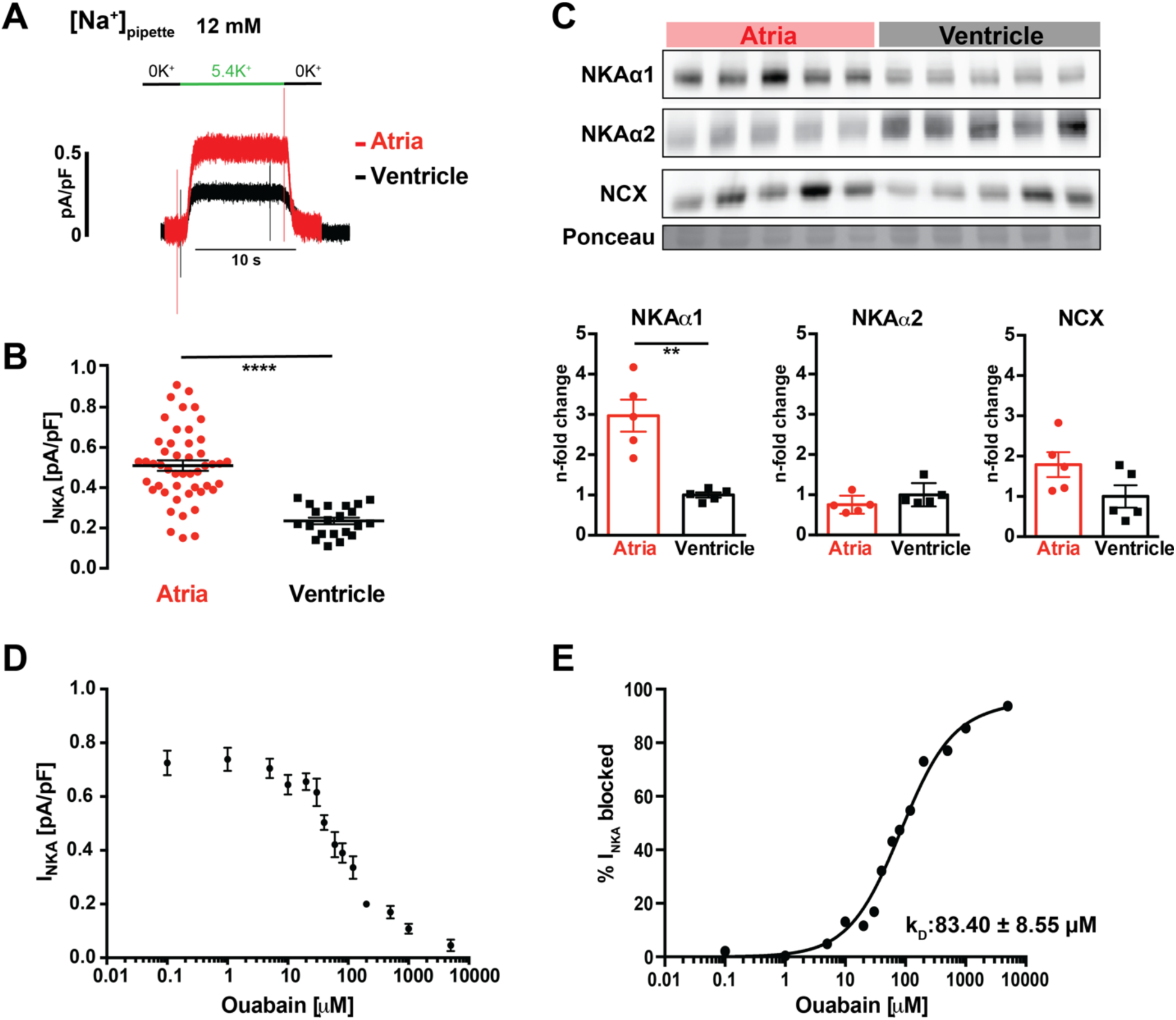
I_NKA_ and NKA protein expression. **A.** Exemplar of I_NKA_ recorded in an atrial (red) and a ventricular (black) myocyte with [Na^+^]_pipette_ = 12 mM. The current is activated by rapidly switching from 0 mM K^+^ (black line) to 5.4 mM K^+^ (green line) in the external solution. **B.** I_NKA_ density in atrial and ventricular murine myocytes (Atria: N = 10 animals, n = 48 cells; ventricle: N = 10 animals, n = 21 cells, **** = p < 0.0001). **C.** Protein expression levels of NKAα1, NKAα2 and NCX in atrial and ventricular tissue normalized to Ponceau. Atrial protein expression levels are expressed as n-fold change over ventricle. **D.** Ouabain-dose response of I_NKA_ in atrial myocytes (N = 10 animals, n = 101 cells). **E.** Fit of data shown in **D**.

### Intracellular [Na^+^]_i_ and [Ca^2+^]_i_ in atrial and ventricular myocytes

Intracellular [Na^+^]_i_ and [Ca^2+^]_i_ were measured in atrial and ventricular myocytes. [Na^+^]_i_ was measured in intact myocytes using the Na^+^-sensitive, ratiometric indicator SBFI as described previously^25,26^. Fig. 5A shows the primary ratiometric data from SBFI measurements in an atrial myocyte along with the calibration information for that cell. Fig. 5B shows the same for a ventricular myocyte. Figs. 5C and 5D show that atrial myocytes have a much lower [Na^+^]_i_ at all stimulation rates (1-3 Hz) and at rest, when compared to ventricular myocytes. [Ca^2+^]_i_ was measured using the Ca^2+^ indicator fura2 in its cell permeant form (Fura2 acetoxymethyl ester, fura2-AM)^27^. A calibration of the fluorescence signal was performed in each myocyte *in situ*. An important advantage of this calibration method is that it corrects for the variability of Fura2 de-esterification between cells by determining the extent of esterification in each individual cell^27^. This information greatly improves the sensitivity of the [Ca^2+^]_i_ quantification. Importantly, the measurements were performed in intact myocytes so that the physiological [Na^+^]_i_, which is significantly different between atrial and ventricular myocytes, remained unperturbed. Fig. 5E shows exemplars of Ca^2+^ transients recorded from murine atrial (red) and ventricular (black) myocytes at different stimulation rates using external field stimulation (0-2 Hz). The diastolic [Ca^2+^]_i_ measured in atrial myocytes (Fig. 5F) is about 30-50 nmol/L lower than the diastolic [Ca^2+^]_i_ in ventricular cardiac myocytes. Similarly, the systolic [Ca^2+^]_i_ in atrial myocytes is lower across all stimulation frequencies compared to ventricular myocytes.

**Figure 5:**
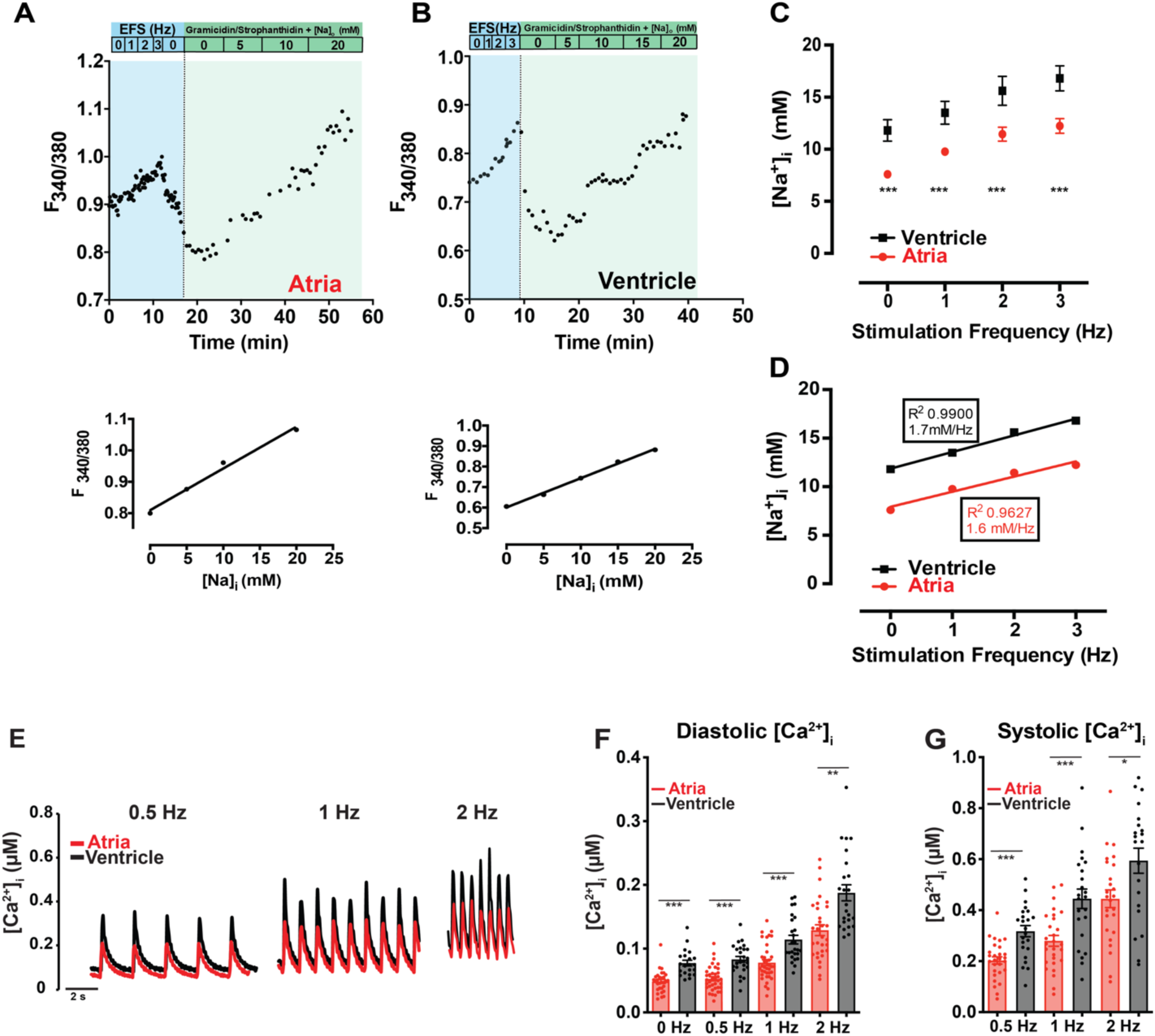
[Na^+^]_i_ and [Ca^2+^]_i_ in murine cardiac myocytes. **A.** Top. Exemplar of SBFI fluorescence (F_340/380_) measured in a murine atrial myocyte during 0-3 Hz external field stimulation (blue) followed by in-situ calibration of [Na^+^]_i_ with increasing [Na^+^]_o_ solutions in the presence of 10 μmol/l gramicidin D and 100 μmol/l strophanthidin (green). Bottom. Determination of [Na^+^]_i_ by linear regression of values measured in top panel. **B.** Exemplar of SBFI fluorescence measurement in a murine ventricular myocyte, as shown in **A**. **C.** Rate dependent [Na^+^]_i_ in atrial and ventricular murine myocytes (Atria: N = 10 animals, n = 16-55 cells; Ventricles: N = 8 animals, n = 16-20 cells). **D.** Regression lines fit to data from **C. E.** Exemplars of whole cell Ca^2+^ transients in atrial and ventricular cardiac myocytes recorded during external field stimulation (0.5-2 Hz). Fura-2 fluorescence (F_340/380_) is calibrated in-situ in each cell. **F.** Rate-dependent diastolic [Ca^2+^]_i_ in atrial and ventricular murine myocytes **G.** Rate-dependent systolic [Ca^2+^]_i_ in atrial and ventricular murine myocytes (Atria: N = 5 animals, n = 19 cells; Ventricles: N = 6 animals, n = 20 cells).

### [Na^+^]_i_ -dependence of NKA activity as measured by I_NKA_

Using the same patch-clamp approach as before (see Fig. 4A) but with [Na^+^]_pipette_ ranging from 5 mmol/L to 100 mmol/L we measured the [Na^+^]_i_-dependence of I_NKA_. Exemplars of I_NKA_ at constant [K^+^] and select [Na^+^]_i_ for atrial and ventricular myocytes are shown in Fig. 6A (red and black traces, respectively). The ß-adrenergic regulation of NKA was examined by using a high concentration of isoproterenol (iso, 1 μmol/L) and measuring its effect on I_NKA_ in atrial and ventricular myocytes (dark red and dark grey traces, respectively, in Fig. 6A) as a function of [Na^+^]_pipette_. The data from these experiments are shown in Fig. 6B and were fit with an equation^28^, which assumes activation by three independent and identical Na^+^-binding sites^28, 29^. The atrial myocyte responsiveness to ß-adrenergic stimulation is huge and occurs at low [Na^+^]_i_. In contrast, the ventricular myocyte responsiveness is relatively weak, occurs at triple the [Na^+^]_i_ and is only significant at a single level of [Na^+^]_i_, 12 mM. The K_1/2_ values for [Na^+^]_i_ in the four curves are shown in Fig. 6C and the (I_NAK_)_max_ values are shown in Fig. 6D. In order to further assess physiological activation pathways of NKA, we measured I_NKA_ after treatment of cells with epinephrine (1 μM, with 12 mM [Na^+^]_pipette_, Fig. 6F). Thus, while the NKA is activated by [Na^+^]_i_ in both cell types, it is much more sensitive to activation by [Na^+^]_i_ in atrial myocytes (K_1/2_ ∼ 3 mM) compared to ventricular myocytes (K_1/2_ ∼ 9 mM). Second, while there is a huge increase of sensitivity of NKA in atrial myocytes when treated with isoproterenol (doubling), there is only a small, albeit significant increase in ventricular myocyte sensitivity to [Na^+^]_i_ when treated with isoproterenol at physiological [Na^+^]_i_.

**Figure 6:**
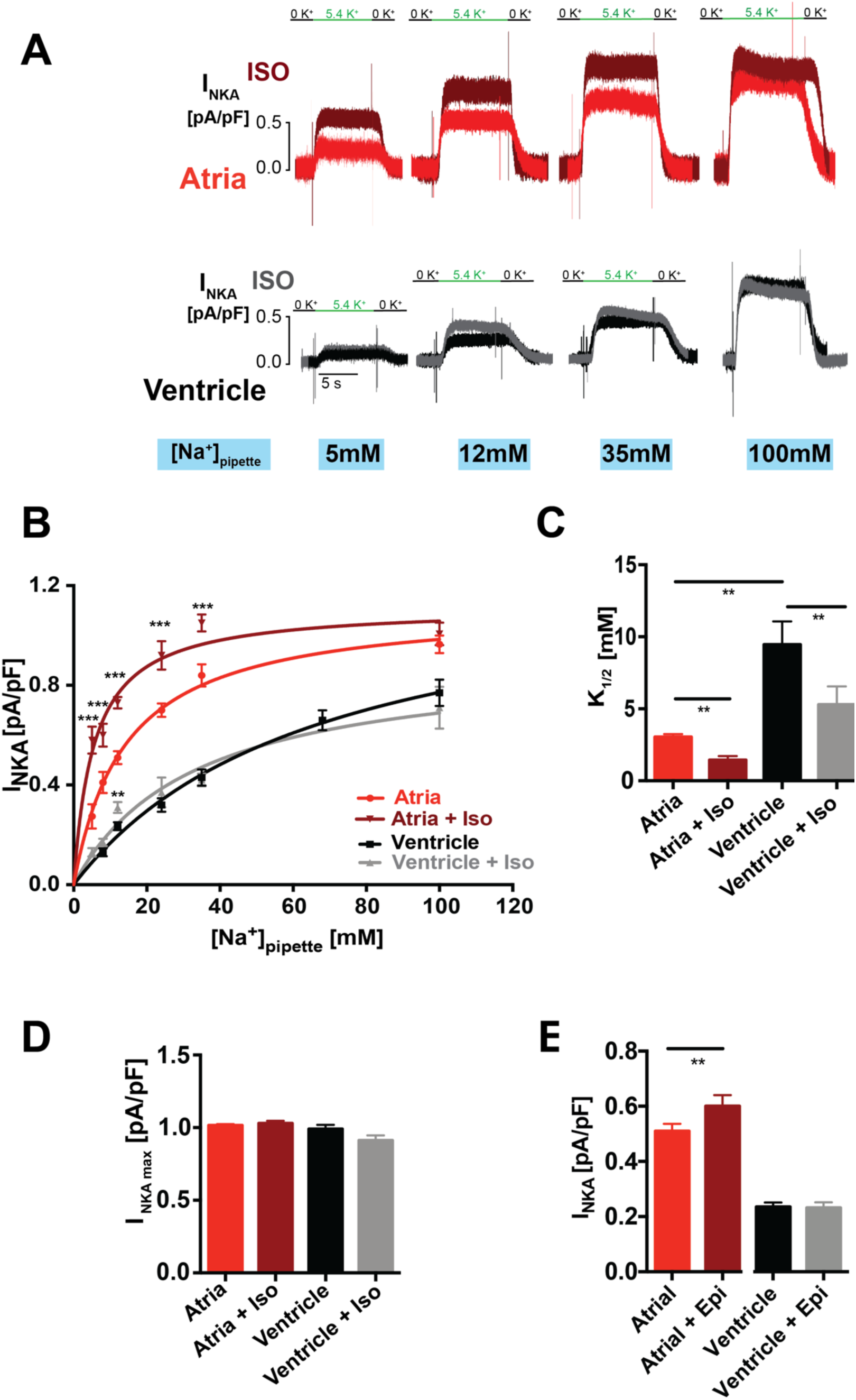
ß-adrenergic regulation of I_NKA_. **A.** Exemplars of I_NKA_ recordings in atrial myocytes with increasing [Na^+^]_pipette_ (red). Traces in dark red show I_NKA_ after treatment with isoproterenol (1 μmol/l). **B.** Exemplars of I_NKA_ from ventricular myocytes, as in **A.** Black traces show I_NKA_ at baseline, grey traces show I_NKA_ after isoproterenol treatment**. C.** [Na^+^]_pipette_-dependent mean I_NKA_ density in atrial and ventricular myocytes with and without isoproterenol (Atria: N = 6 animals, 25 cells; Ventricle: N = 6 animals 18 cells,). **D.** K_1/2_ for [Na^+^]_pipette_ in atrial and ventricular myocytes with and without isoproterenol treatment, derived from fitted data shown in **C**. **E.** I_max_ for atrial and ventricular myocytes with and without isoproterenol treatment, derived from fitted data shown in **C**. **F.** I_NKA_ current density (with [Na^+^]_pipette_ = 12 mM) in atrial and ventricular myocytes with and without epinephrine treatment (1 μmol/l, Atria: N = 4 animals, 12 cells Ventricle: N = 4 animals 14 cells).

### The FXYD1 protein phospholemman, a regulator of NKA function

The NKA is often associated with a small regulatory protein with a FXYD amino acid motif ^30^. In ventricular tissue, FXYD1 or phospholemman (PLM), is associated with NKAα1 and exerts a tonic inhibition of NKA, which is relieved when PLM is phosphorylated (via PKA^31^ and PKC^32^). To investigate the organization of the NKA complex in atrial tissue, co-immunoprecipitation (Co-IP) experiments were carried out in atrial and ventricular tissue. Fig. 7A shows that NKAα1 is associated with PLM in mouse atrial tissue as well as the previously reported association of NKAα1 and PLM in ventricular tissue^33, 34^. PLM is the only FXYD protein that contains phosphorylation sites^35^. Fig. 7B shows protein expression levels for NKAα1 and PLM in atrial and ventricular tissue. Although PLM expression levels are similar in mouse ventricle and atria, the high abundance of NKAα1 in atrial tissue (also shown in Fig. 4C) leads to a much lower PLM/NKAα1 ratio in atria compared to ventricle. When unphosphorylated PLM is associated with NKA, the Na^+^ affinity of NKA decreases 2fold^30^. The high baseline Na^+^ affinity of the NKA in atrial cells (Fig. 6 B, C) is due to a much lower PLM\NKAα1 ratio in atria compared to ventricles. We next examined how systemic *in-vivo* ß-adrenergic activation affects PLM phosphorylation levels. Fig. 7C shows the *in-vivo* protocol with exemplars of mouse ECGs. Sedated mice were injected with isoproterenol (100 mg/kg, i. p.) under constant ECG monitoring, which induced a robust increase in heart rate (Fig. 7C,D). Mice injected with vehicle showed no change in heart rate (Fig. 7D). Data shown in Fig. 7E show Western Blots performed in atrial and ventricular tissue harvested at the end of the isoproterenol protocol depicted in Fig. 7C. Systemic isoproterenol application induced a 5fold and 4fold increase in PLM-ser63 and PLM-ser68 phosphorylation, respectively, over controls (dotted line) in atrial tissue, whereas there was no further increase in PLM phosphorylation in ventricular tissue, suggesting a much higher phosphorylation reserve of PLM in atria compared to ventricles. The large atrial phosphorylation reserve is the underlying mechanism for the much higher increase in atrial Na^+^ affinity of NKA after isoproterenol stimulation shown in Fig. 6 A-C.

**Figure 7:**
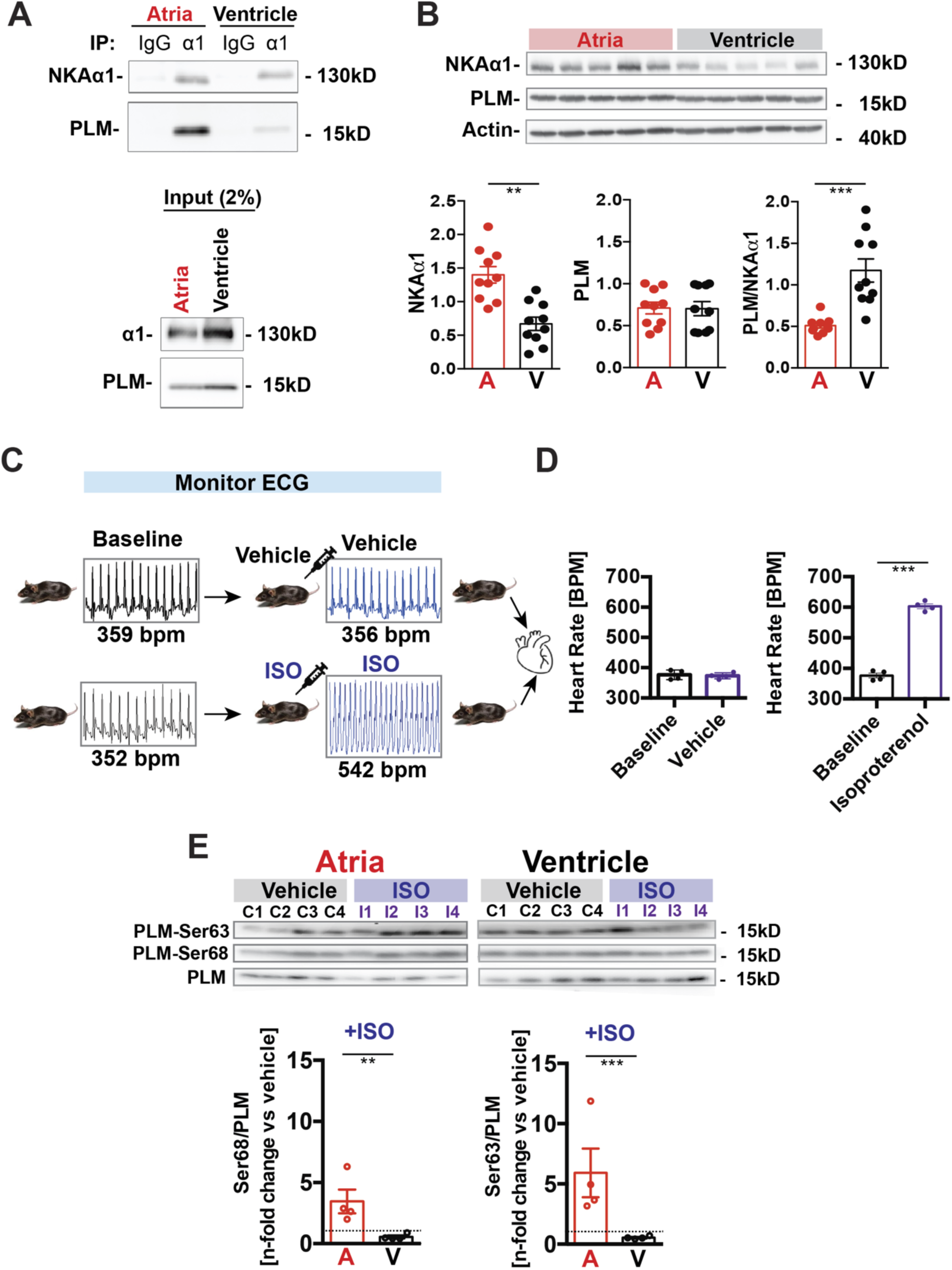
Phospholemman expression and phosphorylation. **A.** Co-immunoprecipitation of NKAα1 with phospholemman (PLM) in atrial and ventricular tissue (N = 15 animals for atria, 5 animals for ventricle). **B.** Western Blot analyses of PLM and NKAα1 protein expression in atrial and ventricular tissue (normalized to actin; n = 5 animals). **C.** Schematic depicting treatment protocol for the evaluation of in-vivo PLM phosphorylation. Isoproterenol was injected i.p. under ECG monitoring. Mice injected with vehicle served as controls. **D.** Mean heart rate before and after either isoproterenol or vehicle injection. **E.** Western Blot analyses of PLM, PLM serine 63 and PLM serine 68 in atrial and ventricular tissue from isoproterenol treated animals and control animals (N = 4 treated animals, n = 4 control animals). Data shown as n-fold change in PLM phosphorylation (PLM-Ser68 and PLM-Ser63) after i. p. injection with isoproterenol compared to controls (animals injected with vehicle).

### Local adrenergic NKA regulation in the NKAα1-Ca_v_1.2 signaling cloud

Quantitative data from STORM super-resolution imaging (Figs. 1 & 2) show that NKAα1 and Ca_v_1.2 clusters share ∼ 80% of their respective area. This means that the accessory and regulatory proteins associated with both ion channels are within range to be effectors in the regulation of both ion channels. This finding is also corroborated by the previously performed proximity proteomics by Marx and colleagues, where association between proteins is assumed to be within ∼40 nm of each other^19^. This means that NKAα1 and Ca_v_1.2 form a signaling nanodomain through shared accessory proteins. To test this hypothesis, we examined whether the high ß-adrenergic sensitivity of atrial I_NKA_, which we found to be mediated by a high PLM phosphorylation reserve (Fig. 6), is locally regulated. Local delivery of protein kinase A (PKA) mediated phosphorylation is often conferred by A-kinase anchoring proteins (AKAPs), which provide scaffolding for PKA close to its phosphorylation targets. Ca_v_1.2 interacts with AKAP15 (also termed AKAP7 or AKAP18) via a leucine zipper-like motif^36^ and with AKAP79/150 in a similar way^37^. Inhibitors of the PKA-AKAP interaction, like the peptide Ht31 and *St*apled *A*KAP *D*isruptor (STAD) peptides have been crucial tools in the investigation of AKAP mediated PKA phosphorylation^38^. Here we used two structurally distinct AKAP disruptor peptides, stHt31 and STAD2, to examine the role of AKAPs in effecting local ß-adrenergic regulation of atrial I_NKA_. We hypothesized that AKAPs within the affinity cluster are local effectors of PKA phosphorylation of PLM. Fig S1A depicts a schematic of the AKAP disruption mechanism. Fig S1B shows an exemplar each of a confocal image of an atrial (i) and a ventricular (ii) myocyte stained with an antibody against the PKA regulatory subunit II ß (PKA RIIß). About 75% of PKA RII is compartmentalized by binding to AKAPs, while PKA RI is cytosolic^39^. Treatment of atrial and ventricular myocytes with the AKAP disruptor stHt31 (50 μmol/l, 20 minutes) shows a reduction in PKA RIIß fluorescence intensity when compared to untreated myocytes (Fig S1B). Fig. 7A shows that in atrial myocytes that were pretreated with either stHt31 (50 μM, 20 minutes) or STAD2 (50-100 nM, 20 minutes) the isoproterenol-induced increase in I_NKA_ was completely blunted. A scrambled stHt31 protein did not change the isoproterenol-induced increase in I_NKA_ (Fig 7A). These data provide clear evidence that the PKA-mediated regulation of I_NKA_ is locally regulated by AKAPs in the NKA-Ca_v_1.2 nanodomain. WB analyses of the regulatory subunits of PKA in atrial versus ventricular tissue show a distinct protein expression pattern with PKA RIIß dominating in atrial and PKA RIIα dominating in ventricular tissue. These data further confirm how the qualitative and quantitative composition of the NKA-CAv1.2 nanodomain underlies tissue specific intracellular Na^+^ homeostasis in cardiac myocytes.

Fig. 7B depicts a schematic of the new cardiac NKA-Ca_v_1.2 signaling cloud. Fig. 7C shows how the novel NKA-Ca_v_1.2 signaling nanodomain integrates into the cardiac dyad.

## Discussion

Our investigation has provided a new understanding of the mechanism by which the excitation-contraction machinery in the heart operates. The core element, the cardiac dyad, traditionally is composed of a set of voltage-sensitive triggering channels, L-type Ca channels (Ca_v_1.2), that are precisely placed in apposition to a cluster of RyR2s channels on the junctional SR. By this means the triggering Ca^2+^ influx is amplified by Ca^2+^-induced Ca^2+^-release (CICR) to produce a Ca^2+^ spark and, when multiple Ca^2+^ sparks are synchronized, a Ca^2+^ transient. We provide strong direct evidence that the core of this nanodomain consists of structural and functional nanodomain of tightly co-clustered NKAα1 and Ca_v_1.2 channels, that has not been conceptualized previously. Our work

This study further provides direct evidence that the organization and composition of the NKA-Ca_v_1.2 nanodomain is tissue specific and affects bulk cellular [Na^+^]_i_ and [Ca^2+^]_i_. Using STORM super-resolution microscopy we show that NKAα1 and Ca_v_1.2 form large superclusters on the cell membranes in atrial myocytes. In addition, these atrial superclusters have a very high ß-adrenergic sensitivity, which is locally regulated by A-kinase anchoring proteins (AKAP). Specifically, in atrial tissue high phosphorylation levels of the NKA regulatory protein phospholemman (PLM) underlie high atrial Na^+^ extrusion rates at baseline and in response to ß-adrenergic activation. These atrial specific features of the signaling cloud form the machinery that results in lower global [Na^+^]_i_ and secondarily lower [Ca^2+^]_i_ in atrial compared to ventricular myocytes.

### The cardiac NKAα1-Ca_v_1.2 nanodomain

This study provides direct evidence that the pore-forming subunit of the dominant cardiac L-type Ca^2+^ channel (Ca_v_1.2) co-localizes with NKAα1, the main cardiac isoform of the α subunit of the Na^+^/K^+^ ATPase to form a structural and functional nanodomain on the sarcolemma and the transverse and axial tubular system in cardiac myocytes. Quantitative 2D super-resolution microscopy provides direct evidence that NKAα1 co-clustering with Ca_v_1.2 results in the formation of overlapping ion channel complexes. The data show that ∼ 80% of the Ca_v_1.2 ion channel cluster area overlaps with the NKAα1 cluster area. Further corroboration of the NKAα1-Ca_v_1.2 nanodomain comes from proximity proteomics data by Marx and colleagues performed in mouse α1_C_-APEX (Ca_v_1.2) cardiac myocytes, where it was shown that NKAα1 was in the top 2 % (1.36%) of peptide abundance within all α1_C_-APEX targets. This represented a ∼ 20fold increase in peptide abundance over non-labeled myocytes (see Fig 2G)^19^. Previous work supports the concept of a cellular signaling nanodomain created by accessory proteins that modulate both NKA and Ca_v_1.2 function. For example, a study in guinea pig cardiac myocytes reported that phospholemman (PLM), the cardiac regulatory protein associated with NKAα, also co-immunoprecipitated with Ca_v_1.2 ^40^. In the same study, the authors showed that PLM modulated the gating of L-type calcium channels^40^. We and others have previously shown that L-type Ca^2+^ current in human atrial myocytes is potently modulated by tyrosine kinases, specifically src^41, 42^. Src has also been shown to form a regulatory complex with NKAα1, where it affects the regulation of Na^+^ extrusion^43^. Previous work by Mohler and colleagues reported that NKA and NCX co-cluster with ankyrin-2, a protein implicated in the membrane targeting of ion channels, on the cardiac SL and TATS^14^. Ankyrin-2 and ankyrin-3 were also found in the top 2 % of peptide abundance within α1_C_-APEX proximity proteomics data (Fig 2G), similar in peptide abundance to NKAα1 from previous work by Marx and colleagues^19^. These data suggest that ankyrins might be involved in targeting the NKAα1-Ca_v_1.2 nanodomain to its location in the dyad.

A-kinase anchoring proteins (AKAPs) are scaffolding proteins that tether PKA to their substrate proteins^44, 45^. Ca_v_1.2 is associated with AKAP 15 and AKAP 150^36^. Here, we have shown that the PKA phosphorylation-mediated increase in I_NKA_ is abolished by the use of two structurally different AKAP-PKA disruptor peptides. These data provide functional evidence that the AKAPs within the Ca_v_1.2-NKAα1 nanodomain mediate the PKA phosphorylation dependent increase in I_NKA_. These data provide further evidence that the Ca_v_1.2-NKAα1 nanodomain acts as a functional signaling nanodomain.

In addition,

### [Na^+^]_i_ and [Ca^2+^]_i_ in cardiac myocytes

Previous work by Despa & Bers, our group and others has shown that [Na^+^]_i_ in quiescent ventricular myocytes from rodents ranges from 10 to 14 mM^8, 9, 25, 46–48^. Despite its importance for cardiac myocyte function and its emerging role in atrial disease (e.g. atrial fibrillation^12, 26^) data on atrial Na^+^ homeostasis are sparse^25^. The present study provides the first comprehensive quantitative assessment of atrial [Na^+^]_i_ and its physiological regulation. Surprisingly, [Na^+^]_i_ in murine atrial myocytes is significantly lower than in ventricular myocytes in quiescent myocytes and over all tested stimulation rates. More specifically, direct comparison of atrial and ventricular myocyte [Na^+^]_i_ shows that [Na^+^]_i_ was a substantial 4 mmol/L lower in quiescent atrial compared to quiescent ventricular murine myocytes (Fig. 5). This difference has important implications for cardiac myocyte function. A decrease of only a few mM in cardiac myocytes’ [Na^+^]_i_ causes the Na^+^/Ca^2+^ exchanger (NCX) to extrude more Ca^2+^ from the cell ^49, 50^, thus lowering the intracellular Ca^2+^ concentration ([Ca^2+^]_i_). Indeed, our results show that systolic and diastolic [Ca^2+^]_i_ is significantly lower in quiescent and stimulated atrial myocytes (Fig 2). Lower [Ca^2+^]_i_ leads to a reduction of the sarcoplasmic reticulum Ca^2+^ load ([Ca^2+^]_SR_) and cardiac inotropy ^49, 50^. The atrial myocytes’ lower [Na^+^]_i_ and [Ca^2+^]_i_ therefore likely contribute to the recently reported much lower fractional shortening in atrial myocytes compared to ventricular myocytes^51^.

Importantly, this study provides direct evidence, for the first time how that the quantitative (NKAα1 and Ca_v_1.2 protein expression levels) and qualitative (PKA subtype expression) tissue-specific composition of an ion channel nanodomain is a major regulator of global differences in [Na^+^]_i_ and [Ca^2+^]_i_ between atrial and ventricular myocytes.

### NKA function and regulation in heart

There are multiple entry pathways for Na^+^ into cardiac myocytes but only one main route of Na^+^ extrusion, the sarcolemmal Na^+^/K^+^ ATPase (NKA). NKA extrudes 3 Na^+^ ions for 2 incoming K^+^ ions while hydrolyzing one molecule of ATP. As the only major Na^+^ extrusion pathway NKA is an essential regulator of [Na^+^]_i_. The NKA protein works as an αβ dimer. The α (catalytic) subunit is the cation transporter and contains the ATP binding site, while the β subunit functions as a chaperone^52^. The primary activator of NKA is Na^+^^53^. Here, Na^+^ affinity of the NKA in atrial and ventricular myocytes was measured using whole-cell voltage clamp experiments and increasing [Na^+^]_pipette_. The results show that atrial myocytes have a more than 3-fold higher affinity for Na^+^ than ventricular myocytes. Previous work has shown that the cation composition in the pipette solution affects the k_1/2_ for Na^+^ when I_NKA_ is measured with reported values for k_1/2_ ranging between 3-20 mM Na^+^ (reviewed here^53^). In this study, under the same experimental conditions k_1/2_ for Na^+^ in atrial myocytes was 3.5 mM vs 10 mM in ventricular myocytes. An important modulator of the cardiac NKA’s Na^+^ affinity is its accessory protein FXYD1, or phospholemman (PLM). Association of the NKA α subunit with PLM decreases the NKA’s Na^+^ affinity 2-fold^30^. It was traditionally assumed, in ventricular myocardium, that one PLM associated with one NKAα subunit^2^. Here we show that the PLM/NKAα ratio is different between atrial and ventricular myocardium. While PLM protein expression levels are similar in atrial and ventricular tissue (Fig 7B), atrial NKAα1 protein expression is 3fold higher than in ventricle (Fig. 7B). The lower PLM/NKAα1 ratio in atrial tissue therefore likely contributes to the higher Na^+^ affinity of atrial NKA. PLM is also the only one of the tissue specific FXYD proteins that can be phosphorylated^54^. PLM was found to be the most prevalent target for PKA phosphorylation in cardiac vesicles^35^. Subsequent work by Bers & Despa and others has shown that PKA and PKC mediated phosphorylation of PLM increases NKA Na^+^ affinity and thus the rate of Na^+^ extrusion^31, 32^. The significantly higher baseline and isoproterenol-sensitive Na^+^ affinity of I_NKA_ in atrial myocytes compared to the benchmark data from ventricular myocytes is likely due to the differences in PLM phosphorylation and the PLM/NKAα1 ratio between atrial and ventricular tissue. PLM phosphorylation levels increase much more substantially in atrial tissue than in ventricular tissue after an *in-vivo* challenge with isoproterenol. Here, differences in PKA isoform expression and potentially phosphatase expression might contribute to the high atrial phosphorylation reserve of PLM.

### Role in cardiac disease

Alterations in [Na^+^]_i_ and [Ca^2+^]_i_ are key features of heart failure^9, 11, 49^ and often underlie cardiac arrhythmogenesis^10–12^. Given the key regulatory function of the NKAα1-Ca_v_1.2 nanodomain for the global intracellular Na^+^ and Ca^2+^ homeostasis, structural and functional changes in the NKAα1-Ca_v_1.2 nanodomain are likely to occur in the course of heart failure and or cardiac arrhythmogenesis. Investigation of the NKAα1-Ca_v_1.2 nanodomain in different forms of heart failure and arrhythmia might identify novel treatment targets for the normalization of [Na^+^]_i_ and [Ca^2+^]_i_ in these conditions.

**Figure 8:**
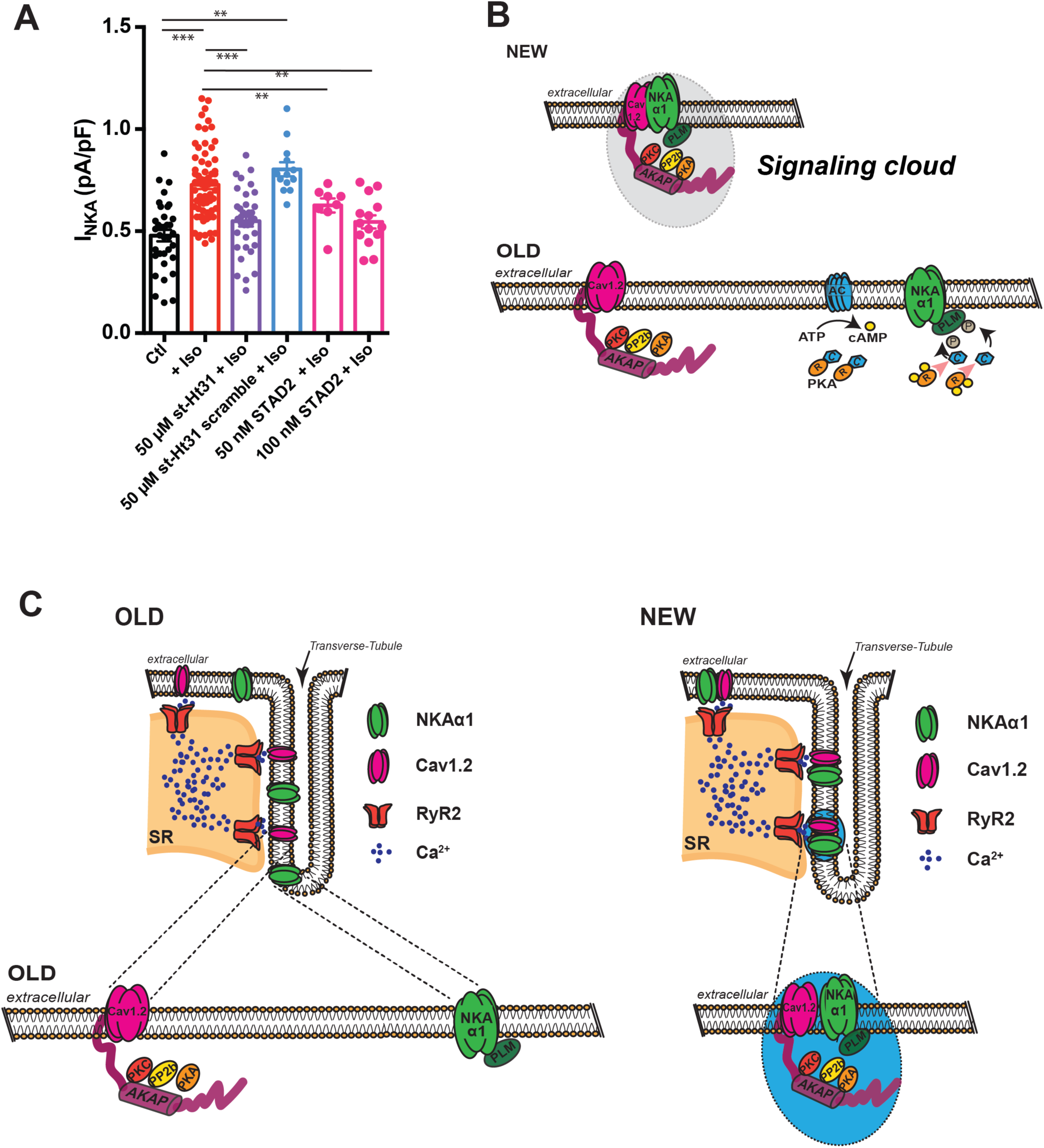
Cardiac NKAα1 and Ca_v_1.2 signaling cloud in atrial myocytes. **A.** I_NKA_ in atrial myocytes with and without 1 μmol/l isoproterenol (data from Fig. 5C) and atrial myocytes pretreated with the AKAP disruptors stHt31 (50 μM, 20 minutes) or STAD2 (50-100 nM, 20 minutes). **B and C.** Schematics of the composition of the novel cardiac Na^+^-Ca^2+^ signaling cloud.

## Non-standard Abbreviations and Acronyms

AKAP: A-kinase anchoring protein
[Ca^2+^]_i_: Intracellular Ca^2+^ concentration
Cav1.2: L-type Ca^2+^ Channel
[Na^+^]_i_: Intracellular Na^+^ concentration
NCX: Na^+^/Ca^2+^ Exchanger
NCLX: Mitochondrial Na^+^/Ca^2+^ Exchanger
NKA: Na^+^/K^+^ ATPase
NKAα1: Na^+^/K^+^ ATPase isoform α1
NKAα2: Na^+^/K^+^ ATPase isoform α2
PKA: Protein Kinase A
PKC: Protein Kinase C
PLM: Phospholemman
PLM-Ser63: Phospholemman Serine 63
PLM-Ser68: Phospholemman Serine 68
PLM-Thr69: Phospholemman Threonine69
RyR2: Ryanodine receptor type 2
SR: Sarcoplasmic Reticulum
TATS: Transverse or axial tubule system
STORM: Stochastic Optical Reconstruction Microscopy

## Sources of Funding

7U19 AI090959 (WJL &LB); Frontiers in Anesthesia Research Award from International Anesthesia Research Society (WJL & LB); R01 HL142290 (WJL &CWW); 5R35GM140822 (WJL); U01 HL116321 (WJL); Special BioMET/UMB Funds (WJL). University of Maryland Claude D. Pepper Center Grant P30 AG028747 (MG), R01 AR071618 (CWW), R01GM129584 (MK).

## Disclosures

None.

## Notes

### Competing Interest Statement

The authors have declared no competing interest.

## References

1. Bers DM and Despa S. Na+ transport in cardiac myocytes; Implications for excitation-contraction coupling. IUBMB Life. 2009;61:215–21.

2. Shattock MJ, Ottolia M, Bers DM, Blaustein MP, Boguslavskyi A, Bossuyt J, Bridge JH, Chen-Izu Y, Clancy CE, Edwards A, Goldhaber J, Kaplan J, Lingrel JB, Pavlovic D, Philipson K, Sipido KR and Xie ZJ. Na+/Ca2+ exchange and Na+/K+-ATPase in the heart. J Physiol. 2015;593:1361–82.

3. Lederer WJ, Niggli E and Hadley RW. Sodium-calcium exchange in excitable cells: fuzzy space. Science. 1990;248:283.

4. Eisner DA and Lederer WJ. Na-Ca exchange: stoichiometry and electrogenicity. Am J Physiol. 1985;248:C189–202.

5. Bers DM, Lederer WJ and Berlin JR. Intracellular Ca transients in rat cardiac myocytes: role of Na-Ca exchange in excitation-contraction coupling. Am J Physiol. 1990;258:C944–54.

6. Crespo LM, Grantham CJ and Cannell MB. Kinetics, stoichiometry and role of the Na-Ca exchange mechanism in isolated cardiac myocytes. Nature. 1990;345:618–21.

7. Blaustein MP and Lederer WJ. Sodium/calcium exchange: its physiological implications. Physiol Rev. 1999;79:763–854.

8. Despa S, Islam MA, Weber CR, Pogwizd SM and Bers DM. Intracellular Na(+) concentration is elevated in heart failure but Na/K pump function is unchanged. Circulation. 2002;105:2543–8.

9. Correll RN, Eder P, Burr AR, Despa S, Davis J, Bers DM and Molkentin JD. Overexpression of the Na+/K+ ATPase alpha2 but not alpha1 isoform attenuates pathological cardiac hypertrophy and remodeling. Circ Res. 2014;114:249–256.

10. Pogwizd SM and Bers DM. Cellular basis of triggered arrhythmias in heart failure. Trends Cardiovasc Med. 2004;14:61–66.

11. Dridi H, Kushnir A, Zalk R, Yuan Q, Melville Z and Marks AR. Intracellular calcium leak in heart failure and atrial fibrillation: a unifying mechanism and therapeutic target. Nat Rev Cardiol. 2020;17:732–747.

12. Sossalla S, Kallmeyer B, Wagner S, Mazur M, Maurer U, Toischer K, Schmitto JD, Seipelt R, Schondube FA, Hasenfuss G, Belardinelli L and Maier LS. Altered Na(+) currents in atrial fibrillation effects of ranolazine on arrhythmias and contractility in human atrial myocardium. Journal of the American College of Cardiology. 2010;55:2330–42.

13. Fischer TH, Herting J, Mason FE, Hartmann N, Watanabe S, Nikolaev VO, Sprenger JU, Fan P, Yao L, Popov AF, Danner BC, Schondube F, Belardinelli L, Hasenfuss G, Maier LS and Sossalla S. Late INa increases diastolic SR-Ca2+-leak in atrial myocardium by activating PKA and CaMKII. Cardiovascular research. 2015;107:184–96.

14. Mohler PJ, Davis JQ and Bennett V. Ankyrin-B coordinates the Na/K ATPase, Na/Ca exchanger, and InsP3 receptor in a cardiac T-tubule/SR microdomain. PLoS Biol. 2005;3:e423.

15. Cunha SR, Hund TJ, Hashemi S, Voigt N, Li N, Wright P, Koval O, Li J, Gudmundsson H, Gumina RJ, Karck M, Schott JJ, Probst V, Le Marec H, Anderson ME, Dobrev D, Wehrens XH and Mohler PJ. Defects in ankyrin-based membrane protein targeting pathways underlie atrial fibrillation. Circulation. 2011;124:1212–22.

16. Popescu I, Galice S, Mohler PJ and Despa S. Elevated local [Ca2+] and CaMKII promote spontaneous Ca2+ release in ankyrin-B-deficient hearts. Cardiovascular research. 2016;111:287–94.

17. Franzini-Armstrong C, Protasi F and Ramesh V. Shape, size, and distribution of Ca(2+) release units and couplons in skeletal and cardiac muscles. Biophys J. 1999;77:1528–1539.

18. Cheng H, Lederer WJ and Cannell MB. Calcium sparks: elementary events underlying excitation-contraction coupling in heart muscle. Science. 1993;262:740–744.

19. Liu G, Papa A, Katchman AN, Zakharov SI, Roybal D, Hennessey JA, Kushner J, Yang L, Chen BX, Kushnir A, Dangas K, Gygi SP, Pitt GS, Colecraft HM, Ben-Johny M, Kalocsay M and Marx SO. Mechanism of adrenergic CaV1.2 stimulation revealed by proximity proteomics. Nature. 2020;577:695–700.

20. James PF, Grupp IL, Grupp G, Woo AL, Askew GR, Croyle ML, Walsh RA and Lingrel JB. Identification of a specific role for the Na,K-ATPase alpha 2 isoform as a regulator of calcium in the heart. Mol Cell. 1999;3:555–63.

21. Despa S and Bers DM. Functional analysis of Na+/K+-ATPase isoform distribution in rat ventricular myocytes. Am J Physiol Cell Physiol. 2007;293:C321–7.

22. Schatzmann HJ. [Cardiac glycosides as inhibitors of active potassium and sodium transport by erythrocyte membrane]. Helv Physiol Pharmacol Acta. 1953;11:346–54.

23. Skou JC. Enzymatic Basis for Active Transport of Na+ and K+ across Cell Membrane. Physiol Rev. 1965;45:596–617.

24. Noel F, Fagoo M and Godfraind T. A comparison of the affinities of rat (Na+ + K+)-ATPase isozymes for cardioactive steroids, role of lactone ring, sugar moiety and KCl concentration. Biochem Pharmacol. 1990;40:2611–6.

25. Garber L, Joca HC, Boyman L, Lederer WJ and Greiser M. Camera-based measurements of intracellular [Na+]_i_ in murine atrial myocytes J Vis Exp. 2022:doi: 10.3791/59600.

26. Avula UMR, Dridi H, Chen BX, Yuan Q, Katchman AN, Reiken SR, Desai AD, Parsons S, Baksh H, Ma E, Dasrat P, Ji R, Lin Y, Sison C, Lederer WJ, Joca HC, Ward CW, Greiser M, Marks AR, Marx SO and Wan EY. Attenuating persistent sodium current-induced atrial myopathy and fibrillation by preventing mitochondrial oxidative stress. JCI Insight. 2021;6.

27. Grynkiewicz G, Poenie M and Tsien RY. A new generation of Ca2+ indicators with greatly improved fluorescence properties. J Biol Chem. 1985;260:3440–50.

28. Gao J, Wang W, Cohen IS and Mathias RT. Transmural gradients in Na/K pump activity and [Na+]I in canine ventricle. Biophys J. 2005;89:1700–9.

29. Gao J, Mathias RT, Cohen IS and Baldo GJ. Two functionally different Na/K pumps in cardiac ventricular myocytes. J Gen Physiol. 1995;106:995–1030.

30. Crambert G, Fuzesi M, Garty H, Karlish S and Geering K. Phospholemman (FXYD1) associates with Na,K-ATPase and regulates its transport properties. Proc Natl Acad Sci U S A. 2002;99:11476–81.

31. Despa S, Bossuyt J, Han F, Ginsburg KS, Jia LG, Kutchai H, Tucker AL and Bers DM. Phospholemman-phosphorylation mediates the beta-adrenergic effects on Na/K pump function in cardiac myocytes. Circ Res. 2005;97:252–9.

32. Han F, Bossuyt J, Despa S, Tucker AL and Bers DM. Phospholemman phosphorylation mediates the protein kinase C-dependent effects on Na+/K+ pump function in cardiac myocytes. Circ Res. 2006;99:1376–83.

33. Bossuyt J, Ai X, Moorman JR, Pogwizd SM and Bers DM. Expression and phosphorylation of the na-pump regulatory subunit phospholemman in heart failure. Circ Res. 2005;97:558–65.

34. Fuller W, Eaton P, Bell JR and Shattock MJ. Ischemia-induced phosphorylation of phospholemman directly activates rat cardiac Na/K-ATPase. FASEB J. 2004;18:197–9.

35. Palmer CJ, Scott BT and Jones LR. Purification and complete sequence determination of the major plasma membrane substrate for cAMP-dependent protein kinase and protein kinase C in myocardium. J Biol Chem. 1991;266:11126–30.

36. Hulme JT, Lin TW, Westenbroek RE, Scheuer T and Catterall WA. Beta-adrenergic regulation requires direct anchoring of PKA to cardiac CaV1.2 channels via a leucine zipper interaction with A kinase-anchoring protein 15. Proc Natl Acad Sci U S A. 2003;100:13093–8.

37. Hall DD, Davare MA, Shi M, Allen ML, Weisenhaus M, McKnight GS and Hell JW. Critical role of cAMP-dependent protein kinase anchoring to the L-type calcium channel Cav1.2 via A-kinase anchor protein 150 in neurons. Biochemistry. 2007;46:1635–46.

38. Kennedy EJ and Scott JD. Selective disruption of the AKAP signaling complexes. Methods Mol Biol. 2015;1294:137–50.

39. Dell’Acqua ML and Scott JD. Protein kinase A anchoring. J Biol Chem. 1997;272:12881–4.

40. Wang X, Gao G, Guo K, Yarotskyy V, Huang C, Elmslie KS and Peterson BZ. Phospholemman modulates the gating of cardiac L-type calcium channels. Biophys J. 2010;98:1149–59.

41. Greiser M, Halaszovich CR, Frechen D, Boknik P, Ravens U, Dobrev D, Lückhoff A and Schotten U. Pharmacological evidence for altered src kinase regulation of I (Ca,L) in patients with chronic atrial fibrillation. Naunyn Schmiedebergs Arch Pharmacol 2007;375:383–392.

42. Schröder F, Klein G, Frank T, Bastein M, Indris S, Karck M, Drexler H and Wollert KC. Src family tyrosine kinases inhibit single L-type: Ca2+ channel activity in human atrial myocytes. Journal of molecular and cellular cardiology. 2004;37:735–745.

43. Tian J, Cai T, Yuan Z, Wang H, Liu L, Haas M, Maksimova E, Huang XY and Xie ZJ. Binding of Src to Na+/K+-ATPase forms a functional signaling complex. Mol Biol Cell. 2006;17:317–26.

44. Gray PC, Scott JD and Catterall WA. Regulation of ion channels by cAMP-dependent protein kinase and A-kinase anchoring proteins. Curr Opin Neurobiol. 1998;8:330–4.

45. Gray PC, Johnson BD, Westenbroek RE, Hays LG, Yates JR, 3rd, Scheuer T, Catterall WA and Murphy BJ. Primary structure and function of an A kinase anchoring protein associated with calcium channels. Neuron. 1998;20:1017–26.

46. Despa S, Islam MA, Pogwizd SM and DM. B. Intracellular [Na+] and Na+ pump rate in rat and rabbit ventricular myocytes. J Physiol. 2002;539:133–43.

47. Baartscheer A, Schumacher CA and Fiolet JW. Small changes of cytosolic sodium in rat ventricular myocytes measured with SBFI in emission ratio mode. Journal of molecular and cellular cardiology. 1997;29:3375–83.

48. Boguslavskyi A, Pavlovic D, Aughton K, Clark JE, Howie J, Fuller W and Shattock MJ. Cardiac hypertrophy in mice expressing unphosphorylatable phospholemman. Cardiovascular research. 2014;104:72–82.

49. Despa S and Bers DM. Na⁺ transport in the normal and failing heart - remember the balance. J Mol Cell Cardiol. 2013;61:2–10.

50. Bers DM. Excitation-Contraction Coupling and Cardiac Contractile Force. 2nd ed. Dordrecht, the Netherlands: Kluwer Academic Publishers; 2001.

51. Nollet EE, Manders EM, Goebel M, Jansen V, Brockmann C, Osinga J, van der Velden J, Helmes M and Kuster DWD. Large-Scale Contractility Measurements Reveal Large Atrioventricular and Subtle Interventricular Differences in Cultured Unloaded Rat Cardiomyocytes. Front Physiol. 2020;11:815.

52. Skou JC and Esmann M. The Na,K-ATPase. J Bioenerg Biomembr. 1992;24:249–61.

53. Glitsch HG. Electrophysiology of the sodium-potassium-ATPase in cardiac cells. Physiol Rev. 2001;81:1791–826.

54. Yap JQ, Seflova J, Sweazey R, Artigas P and Robia SL. FXYD proteins and sodium pump regulatory mechanisms. J Gen Physiol. 2021;153.

