## Supplemental Material for "Na^+^/K^+^ ATPase-Ca_v_1.2 nanodomain differentially regulates intracellular [Na^+^], [Ca^2+^] and local adrenergic signaling in cardiac myocytes"

**Methods**

***Animals***

Male adult C57BL/6 mice (University of Maryland) between 6-8 weeks of age were used in this study.

***Isolation of Atrial and Ventricular Myocytes***

Atrial and ventricular myocytes were isolated using retrograde Langendorff perfusion as previously described ^1, 2^. Animals were anesthetized using isoflurane (4-5 %) inhalation anesthesia administered via a vaporizer. 20 minutes prior to euthanasia mice were injected with an intraperitoneal heparin bolus (1000 U/kg). A thoracotomy was performed, hearts were excised and quickly immersed in ice-cold isolation buffer containing 130 mM NaCl, 5.4 mM KCl, 0.5 mM MgCl2, 0.33 mM NaH_2_PO_4_, 10 mM D-glucose, 10 mM taurine, 25 mM HEPES and 0.5 mM EGTA (pH 7.4, adjusted with NaOH). The aorta was canulated and hearts were then mounted on a Langendorff perfusion system. Hearts were perfused with enzymatic solution containing 1 mg/ml collagenase type II, 0.06 mg/ml protease type XXIII and 0.06 mg/ml trypsin for 5-8 minutes at 37 °C. The atrial chambers were then separated, cut into small tissue strips and subjected to an additional 10 min of enzymatic digestion at 37 °C. Atrial and ventricular myocytes were mechanically dissociated from the tissue by light agitation using a fire polished Pasteur pipette. The cell suspension was then filtered through a nylon mesh (pore size 300 µm) and maintained in modified Tyrode’s solution (in mM: NaCl 130, KCl 5.4, CaCl_2_ 1.8, MgCl_2_ 0.5, NaH_2_PO_4_ 0.33, Glucose 5, HEPES 5 / pH 7.4 adjusted with NaOH) supplemented with 2,3-Butanedione monoxime (BDM,10 mmol/L) and BSA 1 mg/ml. BDM was washed out 30 min prior to experiments by resuspension of cardiac myocytes in Normal Tyrode’s solution. All procedures and protocols involving animal use were approved by the Institutional Animal Care and Use Committee of the University of Maryland School of Medicine (IACUC # 0921015).

***Measurements of [Na^+^]_i_***

Atrial and ventricular [Na^+^]_i_ was measured as described previously ^1, 3^. Freshly isolated myocytes were loaded with the Na^+^ indicator SBFI-AM (10 μM) for 60 minutes at room temperature. After 30 minutes of de-esterification cells were seeded on a laminin coated coverslip. Cells were mounted on an inverted microscope (Nikon Eclipse Ti2) connected to an EMCCD camera (Princeton Instruments Pro EM HS) and a rapid-switching illuminator with a 300 W xenon light source (DG5-plus, Sutter). Cells were continuously perfused with modified Tyrode's solution (in mM: NaCl 130, KCl 5.4, CaCl_2_ 1.8, MgCl_2_ 0.5, NaH_2_PO_4_ 0.33, Glucose 5, HEPES 5 / pH 7.4 adjusted with NaOH). Wide field imaging was achieved with 40X objective (Nikon S Fluor, Oil, 1.30 NA) using two excitation wavelengths (340 nm and 380 nm) by fast switching scanning mirrors and narrow bandwidth excitation filters (340 nm ± 10 nm; 380 nm ± 10 nm). Emission light was collected at 510 ± 40 nm. Data were acquired using Nikon NIS-Elements software. SBFI fluorescence (F_340/380_) was collected after stable baseline recordings had been established for 5 minutes in quiescent cells. Atrial myocytes then underwent external field stimulation (Myopacer, 2 ms, 20 V) and F_340/380_ was recorded after steady-state was established. After completion of measurements calibration of the F_340/380_ signal was performed in situ in each cell. The SBFI loaded myocytes were exposed to five extracellular [Na^+^] ([Na^+^]_o_; 0-20 mM in 5 mM steps) in the presence of 10 µmol/L gramicidin D and 100 µmol/L strophanthidin. The solutions with various [Na^+^]_o_ were prepared by mixing, in different proportions, two solutions of equal ionic strength. One solution contained 145 mM Na^+^ (30 mM NaCl and 115 mM sodium gluconate) and no K^+^, while the other contained 145 mM K^+^ (30 mM KCl and 115 mM potassium gluconate) and was Na^+^ free. Both calibration buffers also contained 10 mM HEPES, 10 mM glucose and 2 mM EGTA. The pH was adjusted to 7.2 with Tris base. Wide field imaging was achieved with a 40X objective (Nikon S Fluor, Oil, 1.30 NA) using two excitation wavelengths (340 nm and 380 nm) by fast switching scanning mirrors and narrow bandwidth excitation filters (340 nm ± 10 nm; 380 nm ± 10 nm). Emission light was collected at 510 ± 40 nm. Data were acquired using Nikon NIS-Elements software. Atrial myocytes underwent external field stimulation (2 ms, 20 V) and F_340/380_ was recorded after steady-state was established. After completion of measurements calibration of the F_340/380_ signal was performed in situ in each cell.

***Measurements of [Ca^2+^]_i_***

[Ca^2+^]_i_ in atrial and ventricular myocytes was measured as described previously. Freshly isolated myocytes were loaded with the Ca^2+^ indicator Fura2-AM (5-10 μM) for 20 min at room temperature. After 20 minutes of de-esterification cells were seeded on a laminin coated coverslip. Cells were mounted on an inverted microscope (Nikon Eclipse Ti2) connected to an EMCCD camera (Princeton Instruments Pro EM HS) and a rapid-switching illuminator with a 300 W xenon light source (DG5-plus, Sutter). Wide field imaging was achieved with 40X objective (Nikon S Fluor, Oil, 1.30 NA) using two excitation wavelengths (340 nm and 380 nm) by fast switching scanning mirrors and narrow bandwidth excitation filters (340 nm ± 10 nm; 380 nm ± 10 nm). Emission light was collected at 510 ± 40 nm. Data were acquired using Nikon NIS-Elements software. Fura2 fluorescence (F_340/380_) was collected after steady-state Ca^2+^ transients were established for 1 minute during external field stimulation. Cells were continuously perfused with modified Tyrode's solution (in mM: NaCl 130, KCl 5.4, CaCl_2_ 1.8, MgCl_2_ 0.5, NaH_2_PO_4_ 0.33, Glucose 5, HEPES 5, pH 7.4 adjusted with NaOH). Ca^2+^ transients were elicited by external field stimulation (Myopacer, 2 ms, 20 V). After completion of measurements calibration of the F_340/380_ signal was performed in situ in each cell. Myocytes were permeabilized with 2 μmol/L ionomycin and immobilized by cytochalasin D (50 μmol/L) and EGTA (1mmol/L). Myocytes were perfused with a 0 and 10 mmol/L Ca^2+^ modified Tyrode's solution (see above) containing ionomycin (2 μmol/L), cytochalasin D (50 μmol/L) and EGTA (1 mmol/L) to determine F_min_ and F_max,_ respectively. The F_340/380_ signal from each cell was then calibrated using the following equation^4, 5^:

$$\left[ Ca^{2^{+}} \right]_{i}: k_{d}* \beta*\frac{F-F_{min}}{F_{max}-F}$$

where F_min_, F_max_ and ß have their usual meaning. k_d_ = 0.25 was obtained as described previously ^6, 7^.

***Measurements of I_NKA_***

The Na^+^/K^+^ ATPase current (I_NKA_) was measured as described elsewhere^8, 9^ with modifications. Isolated atrial and ventricular myocytes were voltage-clamped in the whole cell configuration using electrodes made from borosilicate glass. Electrode resistance when electrodes were filled with internal solution was between 2 - 3 MΩ and series resistance was between 5-15 MΩ. I_NKA_ was recorded using a HEKA patch clamp amplifier. Membrane capacitance and series resistance were calculated from a 5 mV voltage step. After obtaining whole-cell configuration cells were held at -40 mV to inactivate I_Na_. The standard pipette solution contained (in mM): 8 NaCl, 20 KCl, 100 K-aspartate, 20 TEA-Cl, 5 HEPES, 5 Mg-ATP, 5 EGTA. In the pipette solution containing 12-100 mM Na^+^, K-aspartate was replaced by equimolar concentrations of aspartic acid and NaOH (pH 7.2). The external solution contained (in mM):136 NaCl, 2 CsCl, NiCl_2_, 1 BaCl_2_, 1 MgCl_2_, 5 HEPES, 10 glucose, and 5.4 KCl or 0 Tris-Cl; pH 7.4. I_NKA_ was measured at -40 mV as the outward current shift induced by switching from K-free to K-containing external solution.

***Immunocytochemistry***

Freshly isolated atrial and ventricular myocytes were seeded on glass-bottom dishes (Cell Nest, 801001), coated with ECM gel (Sigma, E1270) and allowed to settle for 40 minutes. Cells were fixed with 4% PFA for 20 minutes at room temperature and then washed with PBS 3 x 5 minutes prior to 24 h of incubation (4^o^C) with blocking buffer (50% Fish Serum Blocking Buffer (Thermo) and 50% PBS). This was followed by overnight incubation (4^o^C) with primary antibodies against Ca_v_1.2, NKAα1 and Caveolin 3 in blocking buffer. Cells were then washed 3 x 15 minutes with wash buffer (PBS, 0.2% BSA, 0.05% Triton X (v/v)) and then incubated with appropriate Alexa Fluor™ conjugated secondary antibodies (1:200 in blocking buffer) for conventional confocal microscopy or with Alexa Fluor™ 647 and CF 568 secondary antibodies for STORM super-resolution imaging for 90 minutes at room temperature. Excess secondary antibodies were removed by washing 3 x 15 minutes with wash buffer followed by a 3 x 5 minutes wash with PBS.

***Direct Stochastic Optical Reconstruction Microscopy (dSTORM) imaging.***

Prior to image acquisition glucose oxidase imaging buffer was added to the cell dish. Images were acquired on a Nikon Ti2-E inverted microscope with a 100 X NA1.35 silicon oil-immersion TIRF objective (Nikon), and an electron multiplying CCD camera (Princeton Instruments). Cells were pre-bleached for approximately 5 s to push a large proportion of fluorophores into the dark state. Image sequences (20000 cycles per color) were then acquired and analyzed using Nikon Elements software.

***Tissue lysates, immunoprecipitation, and Western blot:***

For tissue lysates, freshly excised ventricle and atria obtained from C57BL/6 mice were cut into small pieces in ice-cold PBS supplemented with protease inhibitors (PBS-PI; cOmplete Mini EDTA-free; ROCHE). Samples were moved to 1.5 ml Eppendorf tubes suspended in PBS-PI and centrifuged at 20,000 x g at 4°C for 10 min. The pellets were resuspended in 2 x SDS sample buffer (Thermo Fisher Scientific) supplemented with 5% β-mercaptoethanol (Millipore) and homogenized using Eppendorf tube fitting pestles (SP SCIENCEWARE) until the suspension appeared homogenous. Samples were incubated at 100°C for 10 min and then centrifuged 20,000 x g at 4°C for 5 min. Supernatants were used for SDS PAGE. Protein concentrations were measured directly in the samples using a NanoDrop 1000 spectrophotometer (Thermo Fisher Scientific). 50 μg of protein per sample was separated on 4-20% gradient Novex Tris-glycine polyacrylamide gels (Thermo Fisher Scientific), and transferred onto polyvinylidene fluoride membranes (Bio-Rad Laboratories). Membranes were blocked in 5% blocking-grade nonfat dry milk (Bio-Rad Laboratories) in PBS-Tween20 and incubated with primary antibodies overnight at 4°C, followed by horseradish peroxidase–conjugated anti-mouse (Roche) or anti-rabbit (Roche) secondary antibodies for 60 min at RT. Blots were developed with Super Signal West Pico ECL (Thermo Fisher Scientific). Blots were imaged using Amersham Imager 600 chemiluminescence imager (GE Healthcare Life Sciences). Antibodies used for Western blotting were: anti-NKAα1 (clone 464.4 Sigma-Aldrich, 1:2000) and anti phospholemman (abcam, ab200204, 1:2000). Densitometric evaluations of protein expression were performed using image ImageJ 1.53k (https://imagej.nih.gov/ij/).

For immunoprecipitation, tissue piece pellets were resuspended in 1 ml of ice-cold IP buffer (20 mM Tris-HCl, pH 7.5, 150 mM NaCl, 1 mM EDTA, 0.5% NP-40, 5 mM *N*-ethylmaleimide, and protease inhibitors). Samples were homogenized using Eppendorf tube fitting pestles (SP SCIENCEWARE) until the suspension appeared homogenous, followed by 60 min incubation at 4°C with gentle rotation. Samples were centrifuged 20,000 x g at 4°C for 45 min, and supernatants containing solubilized material were transferred to new tubes. Proteins were immunoprecipitated using anti-NKAα1 (clone 464.4 Sigma-Aldrich) and anti phospholemman (abcam, ab200204) antibodies overnight at 4°C with gentle rotation, followed by the addition of protein A/G agarose beads (Clontech Takara) for additional 4 hr. Beads were washed 5 times with PI buffer. Proteins were eluted using 2.5% Acetic Acid neutralized with 1M Tris HCL pH 7.5, followed by SDS PAGE analysis.

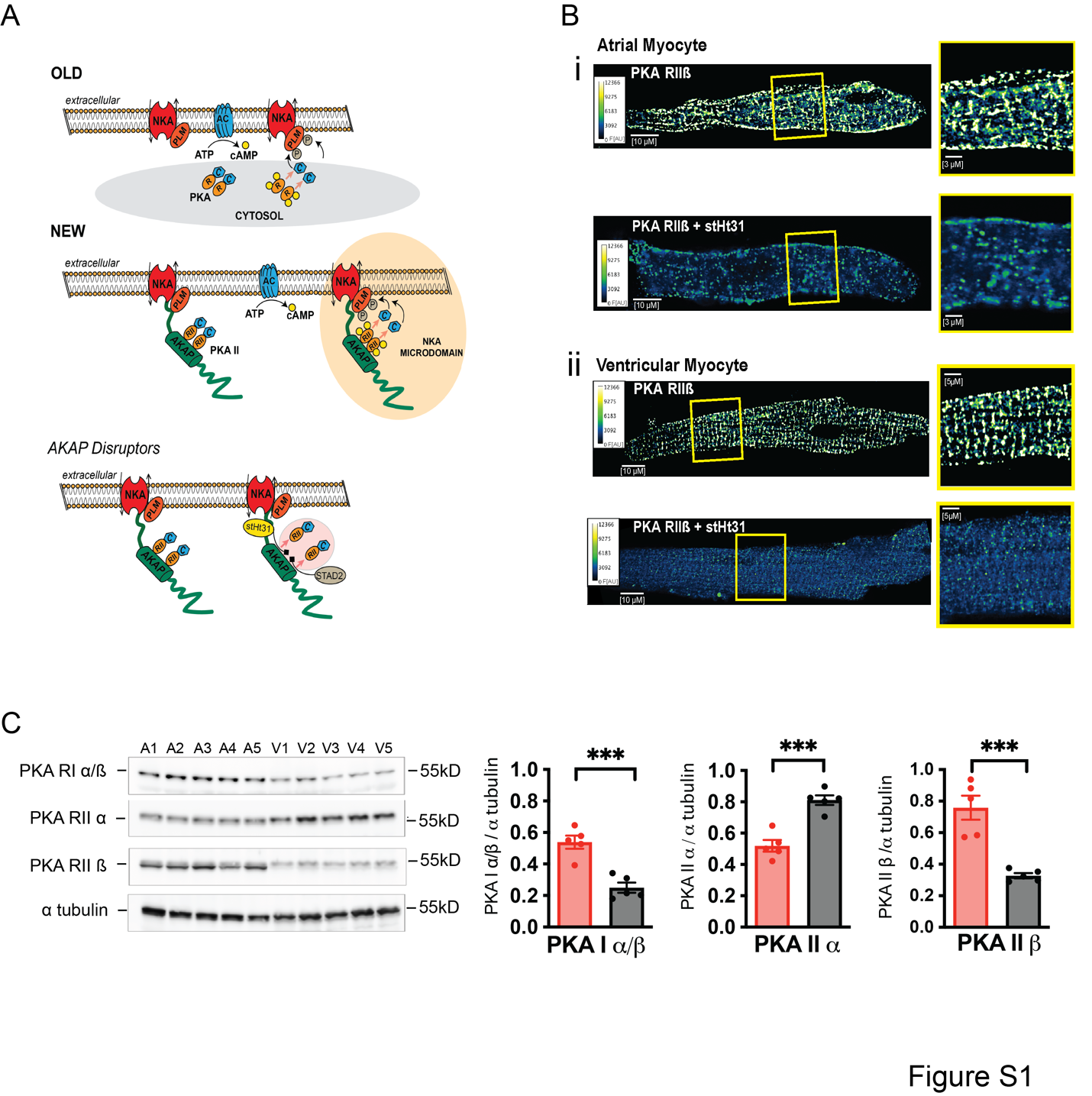

**Figure S1: A.** Schematic depicting the intracellular PKA signaling at the NKA without (old) and with (new) the involvement of local AKAPs. The lowest panel shows the mechanism of action of two AKAP disruptors (stHT31 and STAD2). **B.** Exemplars of an atrial (i) and a ventricular (ii) myocyte stained with an antibody against the regulatory subunit IIβ of PKA with and without prior treatment with the AKAP disruptor stHT31. **C.** Western Blot analyses of PKA regulatory isoforms in mouse atrial and ventricular tissue (normalized to α tubulin).

References

1.	Garber L, Joca HC, Coleman AK, Boyman L, Lederer WJ and Greiser M. Camera-based Measurements of Intracellular [Na+] in Murine Atrial Myocytes. *J Vis Exp*. 2022.

2.	Greiser M, Kerfant BG, Williams GS, Voigt N, Harks E, Dibb KM, Giese A, Meszaros J, Verheule S, Ravens U, Allessie MA, Gammie JS, van der Velden J, Lederer WJ, Dobrev D and Schotten U. Tachycardia-induced silencing of subcellular Ca2+ signaling in atrial myocytes. *The Journal of clinical investigation*. 2014;124:4759-72.

3.	Avula UMR, Dridi H, Chen BX, Yuan Q, Katchman AN, Reiken SR, Desai AD, Parsons S, Baksh H, Ma E, Dasrat P, Ji R, Lin Y, Sison C, Lederer WJ, Joca HC, Ward CW, Greiser M, Marks AR, Marx SO and Wan EY. Attenuating persistent sodium current-induced atrial myopathy and fibrillation by preventing mitochondrial oxidative stress. *JCI Insight*. 2021;6.

4.	Grynkiewicz G, Poenie M and Tsien RY. A new generation of Ca2+ indicators with greatly improved fluorescence properties. *J Biol Chem*. 1985;260:3440-50.

5.	Berlin JR and Konishi M. Ca2+ transients in cardiac myocytes measured with high and low affinity Ca2+ indicators. *Biophys J*. 1993;65:1632-47.

6.	Wescott AP, Kao JPY, Lederer WJ and Boyman L. Voltage-energized Calcium-sensitive ATP Production by Mitochondria. *Nat Metab*. 2019;1:975-984.

7.	Kaplan AD, Boyman L, Ward CW, Lederer WJ and Greiser M. Arrhythmogenic Ca2+ Signaling in a Metabolic Model of HFpEF. *bioRxiv*. 2023:2023.06.21.544411.

8.	Despa S, Islam MA, Pogwizd SM and DM. B. Intracellular [Na+] and Na+ pump rate in rat and rabbit ventricular myocytes. *J Physiol*. 2002;539:133-43.

9.	Workman AJ, Kane KA and Rankin AC. Characterisation of the Na, K pump current in atrial cells from patients with and without chronic atrial fibrillation. *Cardiovascular research*. 2003;59:593-602.
